# Single-molecule imaging reveals the kinetics of non-homologous end-joining in living cells

**DOI:** 10.1101/2023.06.22.546088

**Authors:** Mariia Mikhova, Noah J. Goff, Tomáš Janovič, Joshua R. Heyza, Katheryn Meek, Jens C. Schmidt

## Abstract

Non-homologous end joining (NHEJ) is the predominant pathway that repairs DNA double-stranded breaks (DSBs) in vertebrates. However, due to challenges in detecting DSBs in living cells, the repair capacity of the NHEJ pathway is unknown. The DNA termini of many DSBs must be processed to allow ligation while minimizing genetic changes that result from break repair. Emerging models propose that DNA termini are first synapsed ~115Å apart in one of several long-range synaptic complexes before transitioning into a short-range synaptic complex that juxtaposes DNA ends to facilitate ligation. The transition from long-range to short-range synaptic complexes involves both conformational and compositional changes of the NHEJ factors bound to the DNA break. Importantly, it is unclear how NHEJ proceeds *in vivo* because of the challenges involved in analyzing recruitment of NHEJ factors to DSBs over time in living cells. Here, we develop a new approach to study the temporal and compositional dynamics of NHEJ complexes using live cell single-molecule imaging. Our results provide direct evidence for stepwise maturation of the NHEJ complex, pinpoint key regulatory steps in NHEJ progression, and define the overall repair capacity NHEJ in living cells.

## INTRODUCTION

DNA, the blue-print of life, is remarkably labile and can be damaged in a variety of ways. Damage that results in DNA double strand breaks (DNA breaks on both strands, DSBs) is the most toxic genomic lesion ^1–3^. DSBs that are left unrepaired or repaired incorrectly can result in mutations or loss of genetic information due to chromosomal rearrangements, and, in some cases, cell death. Two major DNA double-strand break repair pathways exist in most organisms ^1–3^. Non-homologous end-joining (NHEJ) directly ligates DNA ends after limited end-processing to render DNA ends chemically compatible for ligation ^2^. NHEJ is active throughout the cell cycle and is estimated to repair 80% of all spontaneous DSBs in human cells ^4^. Homologous recombination (HR) utilizes a sister chromatid to serve as a template DNA to repair DSBs ^5^. Due to the requirement for a sister chromatid, HR functions only in the S and G2 phases of the cell cycle. In addition to engaging repair pathways, DSBs also trigger cell cycle checkpoints that temporarily or permanently halt proliferation or lead to apoptosis of the damaged cell, which is frequently taken advantage of by cancer therapeutics that induce DNA damage ^6^. To understand how human cells respond to DNA damage it is critical to define the repair capacity of the different DNA repair pathways, i.e. the number of DNA lesions that can be repaired per unit time. However, determining the capacity of DNA repair pathways has been challenging, since it requires tracking DNA breaks over time in living cells.

The core NHEJ factors include Ku70, Ku80, DNA-PKcs, XLF, XRCC4 and Ligase 4 ^2,7^. The highly abundant heterodimer of Ku70 and Ku80 (Ku70/80) forms a ring structure that binds double-stranded DNA ends with a high affinity ^8^. Ku70/80 serves as the regulatory subunit of the DNA-dependent protein kinase (DNA-PK), which is activated when Ku70/80 binds DNA ends, resulting in recruitment of DNA-PKcs, the catalytic subunit of DNA-PK ^9^. Ku70/80, and to a lesser extent DNA-PKcs, serve as a scaffold to target the other core NHEJ factors and numerous end-processing enzymes, many of which possess Ku-binding motifs that facilitate assembly of various NHEJ complexes ^7,8,10^.

In recent years, cryogenic electron microscopy (Cryo-EM), and *in vitro* single-molecule imaging studies have significantly advanced our understanding of the assembly mechanism underlying the recruitment of the core NHEJ factors to facilitate DSB repair ^11–19^. Single-molecule Förster resonance energy transfer (FRET) studies suggest that after Ku70/80 binds DNA ends and recruits DNA-PKcs, the DNA ends are synapsed, but are positioned more than 100 Å apart in a long-range synaptic complex ^16,17^. This long-range complex can progress into a short-range synaptic complex where the ends are in sufficiently close proximity to facilitate ligation. This transition requires Ku70/80, the catalytic activity of DNA-PKcs, and components ofthe ligase complex (Lig4, XRCC4, and XLF) ^17^. Recently a plethora of new cryo-EM studies has provided support for this model and surprisingly defined two types of long-range complexes. One long-range complex can form with only Ku70/80 and DNA-PKcs (but can also include XLF, XRCC4, and Ligase4); the other long-range complex is dependent on XLF and has only been observed in complexes that also include XRCC4 and Ligase 4 ^11–15^. Follow-up mutational analysis suggested that the different long-range complexes facilitate distinct end-processing reactions that are required for the repair of DSBs with specific DNA end chemistries ^19^. However, there is currently no direct evidence that these complexes exist in living cells. Moreover, a clear understanding of how NHEJ progresses *in vivo* is lacking. A key challenge has been the lack of tools to study the binding of NHEJ factors to chemically well-defined breaks in living cells. Live cell imaging of laser micro-irradiation (LMI) induced DNA damage has been a useful approach to visualize the recruitment of DNA repair factors to spatially and temporally well controlled DNA lesions in real time ^20–22^. However, LMI induces complex DNA lesions that include DSBs, single-strand breaks, and base damage and the relative contribution of the various types of DNA damage to the recruitment of the repair factors is impossible to disentangle ^23–25^. In addition, because only two to four copies of each NHEJ factor are expected to be recruited to a single DSB, the accumulation of NHEJ factors at DNA lesions is difficult to detect in living cells.

In this study, we address these challenges by developing a single-molecule imaging method to analyze the recruitment of endogenously HaloTagged NHEJ factors to chemically induced DNA breaks. We define the recruitment dynamics of the core NHEJ factors to DSBs and demonstrate that U2OS cells have the capacity to repair ~1000 DSBs per minute via the NHEJ pathway. Our results reveal that Ku70/80 and DNA-PKcs are recruited to DSBs before the components of the ligase complex (XRCC4, XLF, Lig4), consistent with the formation of a long-range complex that only contains Ku70/80 and DNA-PKcs. In addition, we observe that DNA-PKcs dissociates from DSBs ~10 minutes prior to Ku70/80, XLF, and XRCC4. This provides direct evidence for the presence of DNA-PKcs free, long-lived short-range synaptic complexes in cells, and suggests a subset of breaks require extensive processing prior to ligation. Importantly, the dissociation of DNA-PKcs requires its catalytic activity, and its inhibition prevents the release of all other NHEJ factors analyzed, consistent with an arrest of NHEJ progression. Finally, we analyze the role of Ligase 4 in NHEJ complex maturation. In the absence of Ligase 4, the long-range synaptic complex containing XLF and the XRCC4-Ligase 4 fails to form, and Ku70/80 and DNA-PKcs rapidly dissociate from DNA breaks. In addition, consistent with previous in vitro studies, the catalytic activity of Ligase 4 is not required for short-range complex formation, but delays the removal of DNA-PKcs, indicating that the interaction of Ligase 4 with DNA substrate might contribute to the transition to the short-range synaptic complex. Altogether our work provides direct evidence for the existence of NHEJ complexes with distinct compositions in living cells, precisely determines their maturation over time and defines the repair capacity of the non-homologous end joining pathway in cancer cells.

## RESULTS

### Generation of functional, endogenously HaloTagged NHEJ factors

To facilitate live cell single-molecule imaging of NHEJ factors, CRISPR-Cas9 and homology-directed repair were used to insert the sequence encoding a 3xFLAG-HaloTag (Halo) at the N-terminus of the endogenous *Ku70*, *DNA-PKcs* (generated in our previous work ^26^), *XLF*, and *XRCC4* loci in U2OS cells (Fig. 1A). For each gene, all alleles were edited, the insertions were verified using Sanger sequencing, and exclusive expression of the tagged protein was confirmed using western blot (Fig. 1B). To assess subcellular localization of the HaloTagged NHEJ factors, cells were labeled with a fluorescent HaloTag ligand (JFX650), and the tagged proteins were visualized using widefield fluorescence microscopy. In the presence or absence of the DSB-inducing drug zeocin, Halo-Ku70, Halo-DNA-PKcs, and Halo-XRCC4 localized exclusively to the nucleus (Fig. 1C). In contrast, Halo-XLF was detected in both the cytoplasm and the nucleus, consistent with previous observations by others ^27^. To further confirm the cytoplasmic localization of XLF, we performed cellular fractionations of parental U2OS and the Halo-XLF cell line. Both wildtype XLF and Halo-XLF are present in cytoplasm and nucleus (Fig. S1A), demonstrating that XLF cellular localization is not limited to the nucleus in U20S cells. Notably, Halo-DNA-PKcs, Halo-XLF, and Halo-XRCC4 were excluded from nucleoli (Fig. 1C). In contrast, in the absence of DNA damage, Halo-Ku70 was present in both the nucleoplasm and enriched in nucleoli (Fig. 1C). However, in the presence of Zeocin, Halo-Ku70 appeared to be evicted from nucleoli (Fig. 1C), consistent with observations by others ^28^. Importantly, none of the NHEJ factors formed DNA repair foci after induction of DSBs using zeocin, while Halo-Rif1 repair foci were readily detectable after DSBs induction, as expected ^29^. To assess whether the HaloTag disrupts cellular function of the tagged NHEJ factors, we carried out clonogenic survival assays. Zeocin resistance in the four HaloTagged NHEJ factor cell lines was comparable to parental U2OS cells, while DNA-PKcs knock-out cells were highly sensitive to Zeocin (Fig. 1D, Fig. S1B). We conclude that the N-terminal fusion of the 3xFLAG-HaloTag does not significantly alter the DNA repair function of Ku70, DNA-PKcs, XRCC4, or XLF.

**Figure 1.**
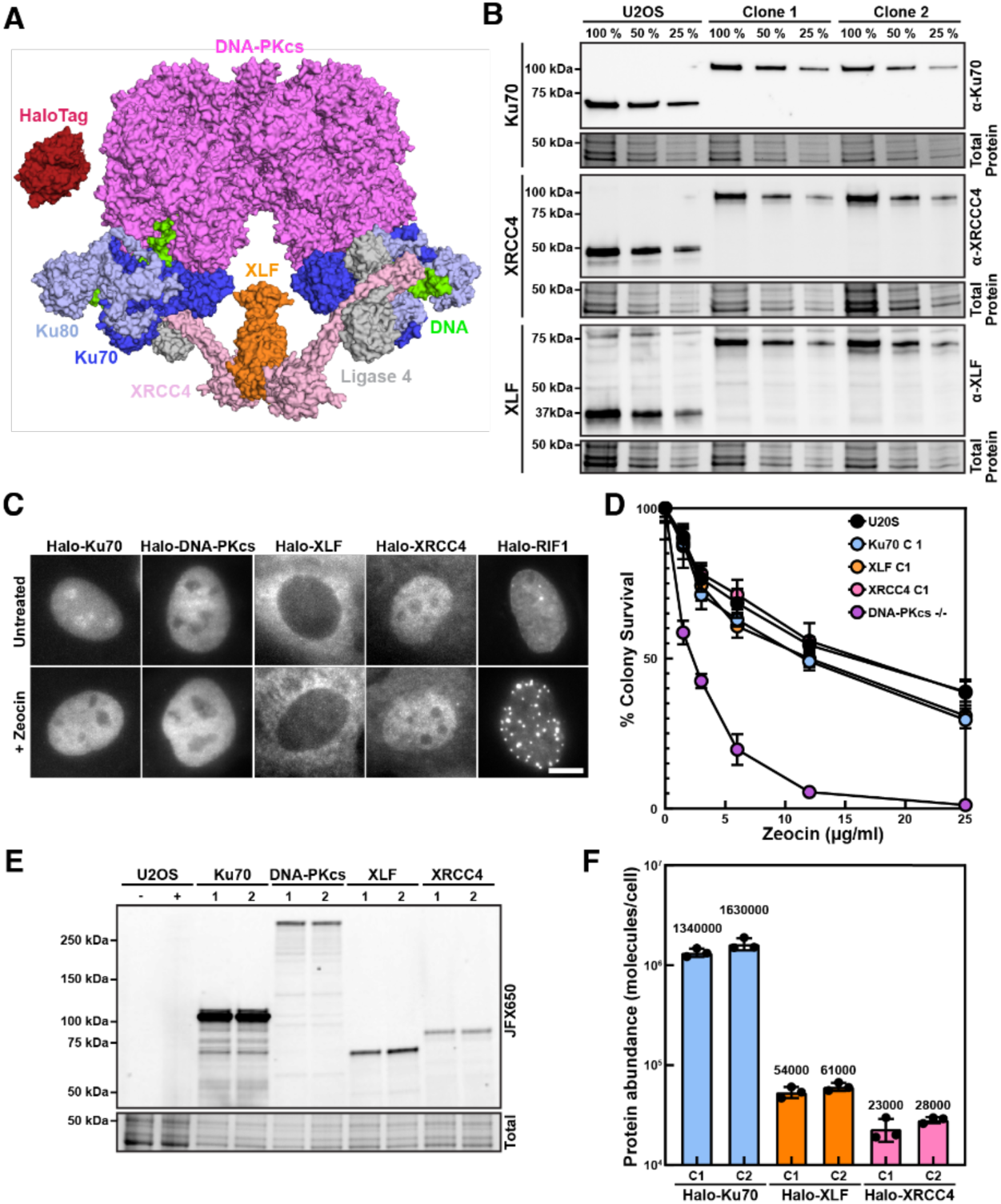
Generation and validation of HaloTagged core NHEJ proteins in U20S cells. **(A)** Graphical depiction of the long-range XLF-mediated NHEJ synaptic complex and HaloTag (based on PDB: 7NFC and 6U32) ^13,44^. **(B)** Western blots of U2OS cells expressing HaloTagged NHEJ factors and parental U2OS cells probed with antibodies against Ku70 (top), XRCC4 (middle), and XLF (bottom). **(C)** Representative images of living cells expressing HaloTagged NHEJ factors and Halo-Rif1 labeled with JFX650 HaloTag ligand in the presence or absence of zeocin (Scale bar = 10 um). **(D)** Clonogenic survival assay of U2OS cells expressing HaloTagged NHEJ factors, parental U2OS cells, and U2OS cells with DNA-PKcs knocked out after challenge with Zeocin (N = 4, 3 technical replicates per biological replicate, Mean ± S.D.). **(E)** Fluorescence imaging of 2 clones of each HaloTagged NHEJ factor labeled with JF650 HaloTag ligand and separated on an SDS-PAGE gel. **(F)** Quantification of the total protein abundance based on recombinant 3X FLAG-HaloTag labeled with JF646 and cell lysates from a specific number of U2OS using in-gel fluorescence intensity values after applying a correction factor from western blots (see Fig. S1C-D, N = 3, Mean ± S.D.).

### Absolute cellular abundance of Ku70, XLF and XRCC4

To determine the absolute cellular abundance of Ku70, XLF, and XRCC4 we compared the fluorescence intensity of labeled Halo-Ku70 and Halo-XRCC4 bands on SDS-PAGE gels to that of a standard containing a known quantity of recombinant 3xFLAG-HaloTag protein labeled with the same fluorophore and a cell lysate from a specified number of cells (Fig. S1C-D). The absolute abundance of Halo-XLF was determined by comparing the fluorescence intensity of Halo-XLF to that of Halo-XRCC4 normalized to total protein loaded (Fig. 1E). To calculate the absolute abundance of endogenous, untagged proteins the absolute abundance of the HaloTagged variants was multiplied by the relative expression of the HaloTagged proteins compared to their untagged counterparts determined by western blot (Fig. 1B). As previously described, the expression level of Ku70 is very high (1.3 - 1.6×10^6^ molecules per cell) ^30^.The expression levels of XLF (54 - 61×10^3^ molecules per cell) and XRCC4 (23 - 28×10^3^ molecules per cell) were substantially lower. In previous work we determined that DNA-PKcs is present at 1.20×10^5^ molecules per cell ^26^. We confirmed the absolute protein number of the tagged NHEJ factors by using a second fluorescent ligand for the HaloTag (JF657, Fig. S1E). Together these observations demonstrate that the abundance of the core NHEJ factors spans two orders of magnitude, with Ku70 being present in a 10-fold excess compared to DNA-PKcs, which in turn is 2-5 fold more abundance than XRCC4 and XLF.

### Stepwise recruitment of NHEJ factors to laser micro-irradiatation induced DNA lesions

To analyze the recruitment of the tagged NHEJ factors to DNA lesions in real time, we performed laser micro-irradiation (LMI) experiments using a 405 nm laser of cells sensitized with Hoechst dye. While LMI induces complex DNA lesions, including DSBs, it provides consistent experimental conditions (pulse length, laser power) and allows analysis of NHEJ factor localization to site of DNA damage with minimal time delay after induction of DNA damage ^20,25^. All tagged NHEJ factors were robustly recruited to LMI-induced damage sites (Fig. 2A-C, Fig. S2A-D, Movie S1-4). Halo-Ku70 and Halo-DNA-PKcs were recruited to damaged sites very rapidly, with a half-maximal accumulation time (t_1/2_) of ~7 and ~17 seconds, respectively (Fig. 3C, Movie S1-2). Halo-XRCC4 and Halo-XLF were recruited with similar kinetics but were delayed (t_1/2_ = 32 and 38 s) compared to Halo-Ku70 and Halo-DNA-PKcs, consistent with previous results (Fig. 2A-C, Movie S3-4) ^31^. Similar to zeocin-induced damage, LMI also induces depletion of Halo-Ku70 from nucleoli (Fig. 2A, Movie S1). These observations demonstrate that the HaloTagged NHEJ factors can bind to DNA lesions and the delayed recruitment of XRCC4 and XLF is consistent with the initial formation of an NHEJ complex composed of only Ku70/80 and DNA-PKcs.

**Figure 2.**
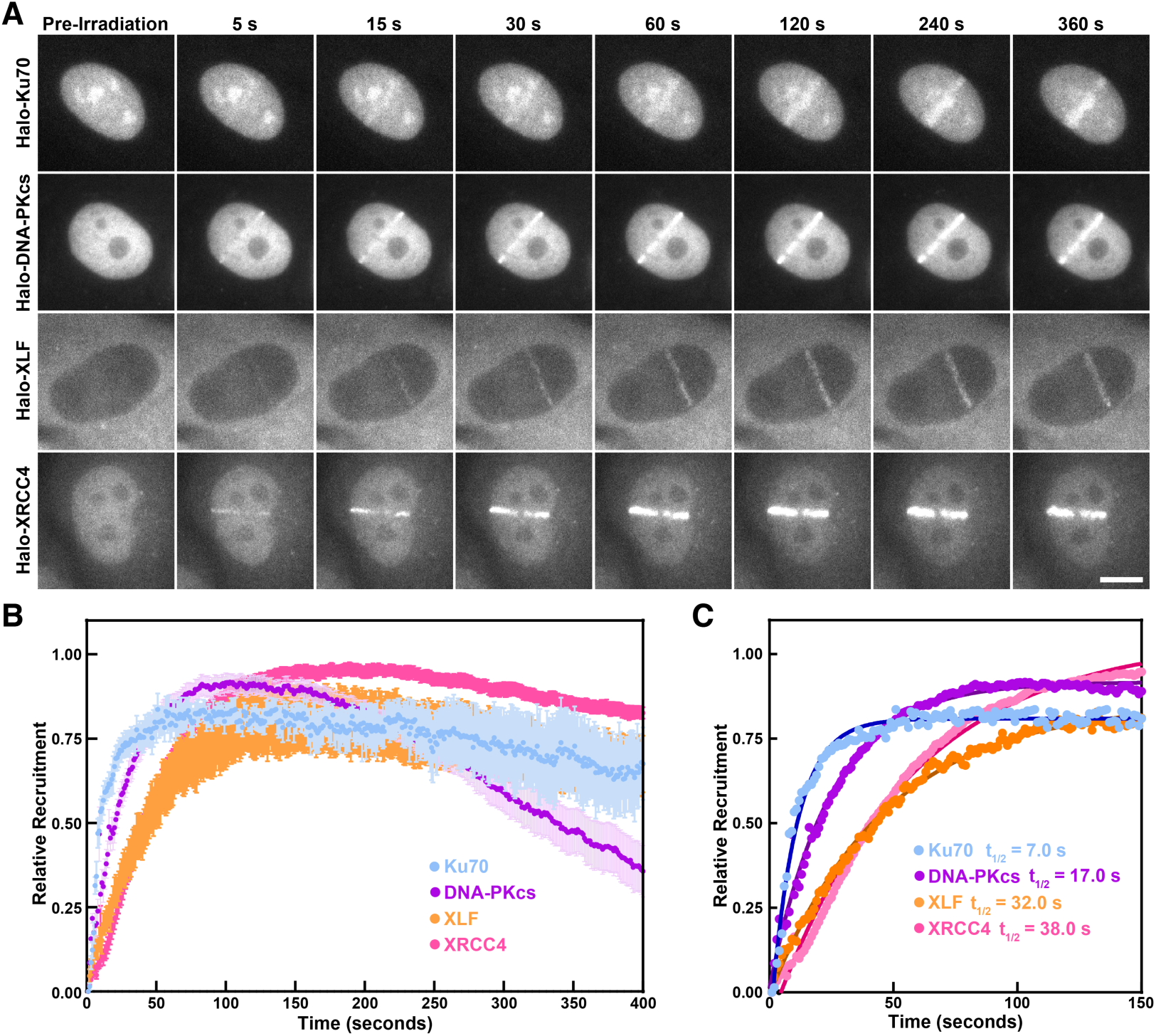
Recruitment kinetics of the core NHEJ factors to complex DNA lesions. **(A)** Representative images of HaloTagged NHEJ factor (JFX650) recruitment to laser-induced DNA lesions over time (Scale bar = 10 μm). **(B)** Normalized recruitment kinetics of HaloTagged DDR proteins to laser-induced DSBs. Data are presented as the average increase in fluorescence post-laser microirradiation normalized to the brightest frame for cell analyzed (N = 10 - 15 individual cells analyzed for each HaloTag cell line, Mean ± S.D.). (**C)** Half-times (t_1/2_) of each protein to laser-induced DSBs determined by nonlinear regression assuming one-phase association (without dissociation).

**Figure 3.**
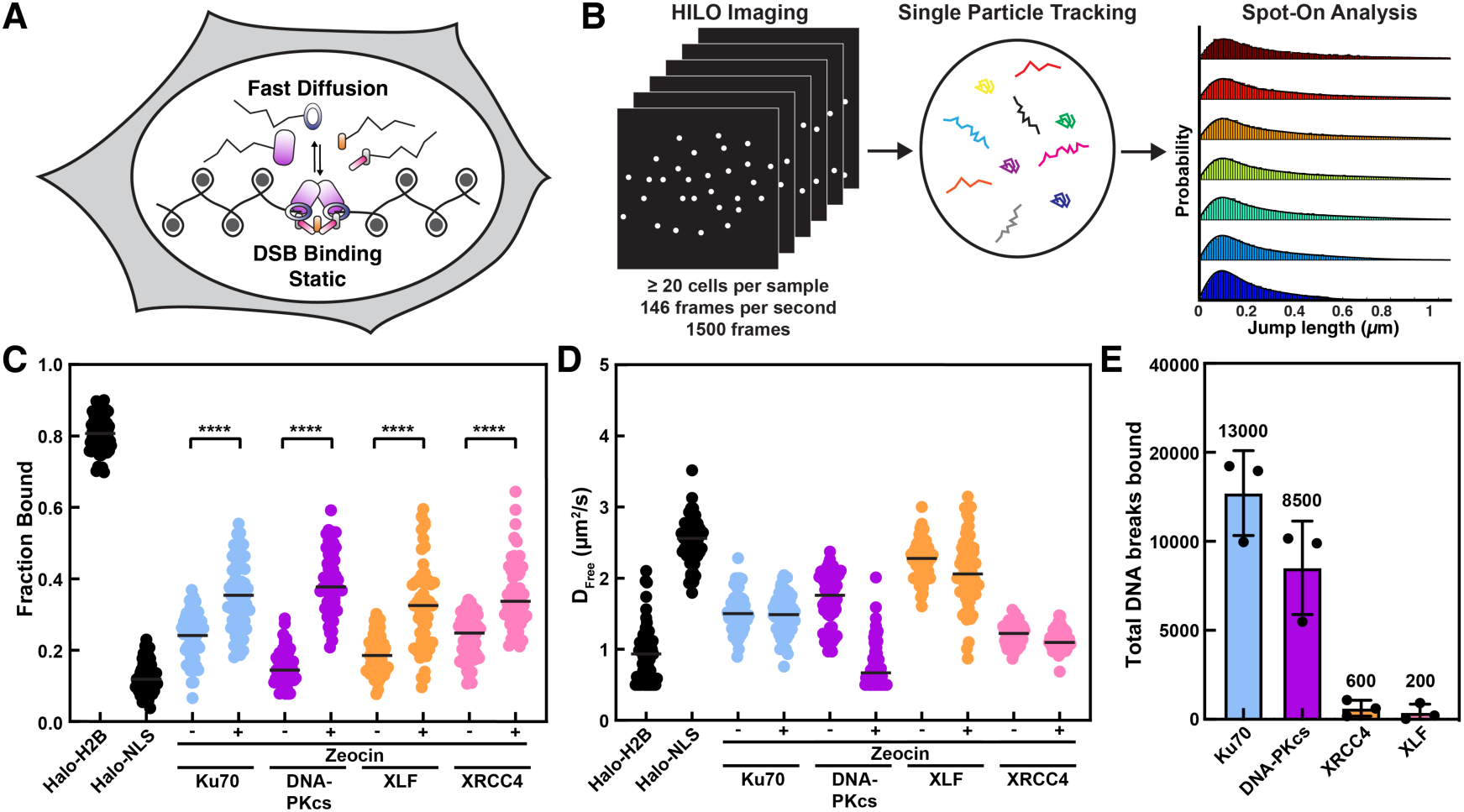
Zeocin-induced DSB formation promotes chromatin association of Halo-tagged NHEJ factors. **(A)** Rational for analyzing DSB association of NHEJ factors using single-molecule imaging. DSB association proteins are static while unbound molecules diffuse rapidly through the nucleus. **(B)** Graphical representation of the workflow used for live-cell single-molecule imaging of core NHEJ proteins. **(C)** Plot of the Fraction Bound for each HaloTag NHEJ protein under untreated conditions and post-zeocin exposure that were analyzed using a two-state model of diffusion. Each data point represents the fraction bound of each protein in an individual cell (N = 3, n ≥ 20 cells for each protein per replicate and condition, Black bar = median). Data were analyzed by two-way ANOVA with Tukey’s posthoc test (**** = p < 0.0001). **(D)** Diffusion coefficients for freely diffusing HaloTagged NHEJ proteins. Each data point represents the D_Free_ calculated from the tracks in an individual cell (N = 3, n ≥ 20 cells for each protein per replicate and condition, Black bar = median). **(E)** Total number of DNA breaks bound be each tagged DNA repair factor after treatment with zeocin (N = 3, Mean ± S.D.).

### Zeocin-induced DSB formation promotes chromatin association of Halo-tagged NHEJ factors

To analyze the diffusion dynamics of individual HaloTagged NHEJ factors, we carried out highly inclined laminated optical sheet (HILO) microscopy of cells sparsely labeled with a fluorescent HaloTag ligand in combination with single-particle tracking (SPT, Movie S5-S9) ^20, 32,33^. Under these conditions only a small fraction of the total cellular proteins was labeled, ranging from 0.01% for Halo-Ku70 to 0.6% for Halo-XLF (Fig. S3A-B). Analysis of single-molecule trajectories using the Spot-On tool allowed us to determine the fraction of static molecules, which we presume include molecules that are directly associated with chromatin, including DSBs (Fig. 3A-B) ^32^. In addition, Spot-On calculates the diffusion coefficients of freely diffusing molecules (D_Free_) and chromatin bound (D_Bound_) molecules (Fig. 3B) ^24^. As controls for freely diffusing and chromatin bound proteins, constructs encoding only the HaloTag fused to three nuclear localization sequences (Halo-NLS) and HaloTag-histone H2B (Halo-H2B) were transfected into parental U2OS cells ^26^. Halo-NLS diffused rapidly through the nucleus (D_Free_ = 2.6 µm^2^/s) and only a small fraction of molecules was static (F_Bound_ = 10%, Fig. 3C-D, Fig. S3D, Movie S5-S6). In contrast and as expected, the majority of Halo-H2B was associated with chromatin (F_Bound_ = 80%, Fig. 3C, Fig. S3D). In the absence of DNA damage, all four HaloTagged NHEJ factors displayed an intermediate level of chromatin bound molecules (F_Bound_ = 20-30%, Fig. 3C, Fig. S3D, Movie S7-S9). Upon DNA damage induction with zeocin the chromatin associated fraction of all tagged NHEJ factors significantly increased by 10-20% (Fig. 3C, Fig. S3D) demonstrating that this single-molecule imaging approach can detect the recruitment of NHEJ factors to DSBs, even though DNA repair foci specific for the core NHEJ factors are generally not detectable by traditional microscopy. The increase in the chromatin bound fraction of DNA-PKcs correlated with the dose of zeocin used (Fig. S3C), consistent with the recruitment of DNA-PKcs to chromatin as a consequence of DSB formation. To confirm that Halo-DNA-PKcs was recruited to chromatin in response to DSB induction in a different cell line, we expressed Halo-DNA-PKcs in CHO-V3 cells (Chinese hamster ovary cells that lack DNA-PKcs). Similar to our observations in U2OS cells, in CHO-V3 cells the chromatin bound fraction of Halo-DNA-PKcs was also increased in response to zeocin treatment (Fig. S3E). The diffusion coefficients of freely diffusing Halo-Ku70, Halo-DNA-PKcs, and Halo-XRCC4 were comparable in the absence of DNA damage (D_Free_ = 1.6-1.9 µm^2^/s, Fig. 3D). Halo-XLF diffused more rapidly through the nucleus (D_Free_ = 2.5 µm^2^/s), indicating that it does not constitutively associate with the other NHEJ factors, in particular XRCC4. The diffusion coefficients of freely diffusing and chromatin bound Halo-Ku70, Halo-XRCC4, and Halo-XLF molecules were not affected by DSBs induction using zeocin (Fig. 3D, Fig. S3F). In contrast, the D_Free_ of Halo-DNA-PKcs was reduced to D_Free_ = 0.7 µm^2^/s after zeocin treatment (Fig. 3D). Similarly, the D_Bound_ of of Halo-DNA-PKcs was also reduced after DSB induction (Fig. S3E). This suggests that DNA-PKcs globally transitions into a different macro-molecular complex after DNA damage induction, or it forms many transient interactions with chromatin that slow down its diffusion dynamics.

In addition to defining the diffusion dynamics of the HaloTagged NHEJ factors, these experiments allowed us to estimate the number of DNA breaks bound by each tagged NHEJ factor per cell. The number of chromatin-bound molecules was determined by multiplying the increase in the static fraction upon zeocin treatment (F_bound zeocin_ – F_bound Untreated_) with the average number of molecules detected per imaging frame for each protein. This number was then divided by the labeling efficiency to account for the small fraction of molecules that are detectable (Fig. S3B). In addition, we multiplied the number by 3 because we only sample a 1 µm thick plane of the cell nucleus, which is 4 µm in height (Fig. S3G, see methods for details) ^34^. Finally, the resulting number was divided by the number of molecules expected to be present at a single DSB (2 for Ku70 and DNA-PKcs, 4 for XRCC4, 2 for XLF, see Fig. 1A). Assuming the increase in the fraction of chromatin-bound molecules after zeocin treatment is exclusively the result of DSB binding, Halo-Ku70 and DNA-PKcs were associated with 13000 and 8500 breaks per cell (Fig. 3E), respectively. Halo-XRCC4 and Halo-XLF only associated with 600 and 200 DSBs per cell (Fig. 3E), respectively. These numbers are consistent with the hierarchy of NHEJ factor recruitment, and the relative abundance of the tagged proteins.

Altogether, these results demonstrate that single-molecule imaging can be used to study the recruitment of NHEJ factors to DSBs in living cells and to estimate the total number of breaks bound by each protein analyzed.

### Kinetics of NHEJ factor recruitment to Calicheamicin induced DSBs

To determine the kinetics of chromatin association of the HaloTagged NHEJ factors, SPT was used to analyze recruitment to calicheamicin induced DSBs; calicheamicin mostly induces DNA breaks with 3’ overhangs, a high percentage of which have 3’ phospho-glycolate adducts ^35,36^. Cultures of cells expressing Halo-Ku70, Halo-DNA-PKcs, and Halo-XRCC4 were treated with various concentrations of calicheamicin for 3 minutes and individual cells were imaged for ~10 seconds each in one-minute intervals, before and after damage induction to determine the fraction of chromatin bound molecules (Fig. 4A, Movie S10, each data point represents a different cell). Prior to treatment with calicheamicin the fraction of chromatin bound molecules remained constant for all NHEJ factors (Fig. 4B, Movie S10). After DNA damage induction the chromatin bound fraction of Halo-Ku70, Halo-DNA-PKcs, and Halo-XRCC4 increased (Fig. 4B, Fig. S4A, Movie S10), consistent with their rapid recruitment to DSBs. Importantly, the fraction bound of Halo-DNA-PKcs cells was not increased after treatment with DMSO (Fig. S4B). In the case of Halo-Ku70 the maximal fraction bound increased as the concentration of calicheamicin was increased, and gradually decreased over time, consistent with NHEJ repairing the DSBs induced by the drug treatment (Fig. 4B,C, Fig. S4A). The time frame for which the fraction bound of Halo-Ku70 remained elevated was elongated with increasing calicheamicin concentration, indicating that the total DSB repair time was increased (Fig. 4B-C, Fig. S4C). In contrast, for Halo-DNA-PKcs and Halo-XRCC4 only small changes in the fraction of chromatin bound molecules were observed as the number of breaks was increased (Fig. 4B, Fig. S4A). In addition, the fraction of chromatin bound Halo-DNA-PKcs and Halo-XRCC4 remained constant over an extended period of time before rapidly dropping (Fig. 4B-C, Fig. S4A,C), rather than gradually decreasing as observed for Halo-Ku70 (Fig. 4B). The duration of this plateau was extended with increasing calicheamicin concentration for both Halo-DNA-PKcs and Halo-XRCC4 (Fig. 4C).

**Figure 4.**
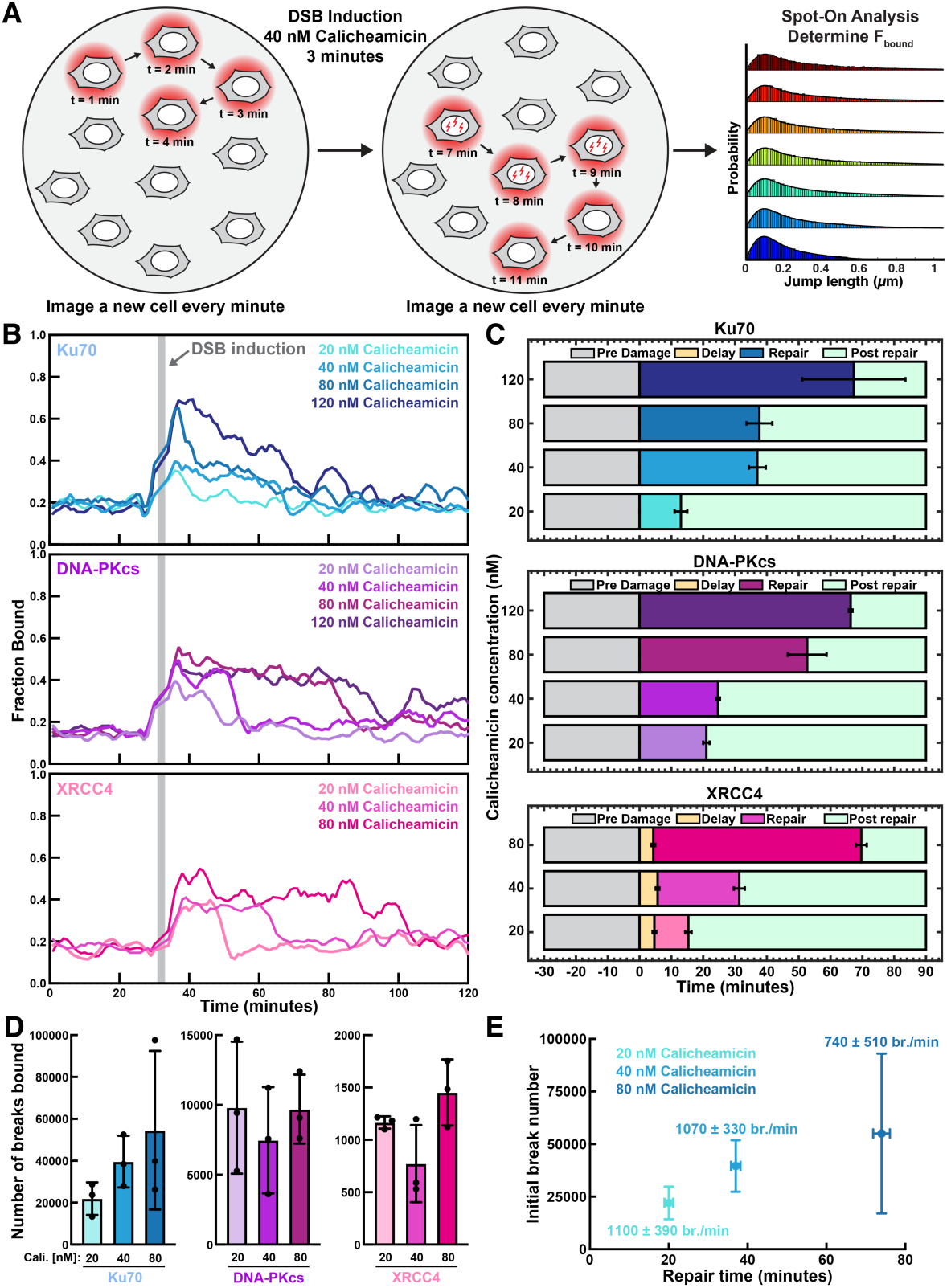
Time course analysis of the recruitment of Halo-tagged NHEJ factors to DNA breaks. **(A)** Experimental approach to analyze the chromatin binding of the tagged DNA repair factors over time. **(B)** Plot of the static fraction of Halo-Ku70, Halo-DNA-PKcs, and Halo-XRCC4 before and after DNA damage induction (shaded area) with various concentrations of calicheamicin for 3 minutes. Graphs represent the rolling average over three consecutive timepoints (for individual graphs with experimental error see Figure S4A). **(C)** Quantification of the timing of the recruitment of Halo-Ku70, Halo-DNA-PKcs, and Halo-XRCC4 after DNA damage induction (N = 3, Mean ± S.D.). **(D)** Quantification of the number of break bound Halo-Ku70 (pooled data for 5 time points after DNA damage induction), Halo-DNA-PKcs (pooled data for 10 time points after DNA damage induction), and Halo-XRCC4 (pooled data for 10 time points after chromatin recruitment) molecules after DNA damage induction with various concentrations of calicheamicin (N = 3, Mean ± S.D.). **(E)** Quantification of the DNA repair rate by using static Halo-Ku70 as a marker for the initial number of DNA breaks (Fig. 4D) and the dissociation of XRCC4 (Fig. 4B) to mark the completion of DNA repair by NHEJ (N = 3, Mean ± S.D.)

To determine the total number of DNA breaks bound by the tagged NHEJ factors, we carried out the same calculations outlined above for zeocin treated cells, averaging the first 5 cells after calicheamicin treatment for Halo-Ku70 or the first 10 cells after DNA recruitment was observed for Halo-DNA-PKcs and Halo-XRCC4. The total number of breaks bound by Halo-Ku70 was elevated from 22000 to 55000 breaks per cell with increasing concentrations of calicheamicin (Fig. 4D). In contrast, Halo-DNA-PKcs and Halo-XRCC4 bound to ~10000 and ~1000 breaks per cell for all calicheamicin concentrations used, respectively (Fig. 4D). These observations suggest that the number of breaks induced in this experiment exceeds the capacity of DNA-PKcs and XRCC4 and these factors are functioning at maximum speed, conceptually similar to an enzyme in the presence of large excess of substrate (Fig. S4D). In contrast, the chromatin recruitment of Ku70 appears not to be saturated, consistent with Ku70 being present in at least a 10-fold excess compared to DNA-PKcs and XRCC4. It is important to note that we are not measuring the repair timing of individual DNA breaks in this experiment since we are only imaging each cell for 10 seconds. Instead, these experiments report on the rate with which the aggregate of all DSBs are repaired in the cell population. In total, these results are consistent with Ku70 rapidly binding most, if not all, DSBs that are induced, which exceeds the total number of cellular DNA-PKcs and XRCC4 molecules (Fig. S4D). Thus, in order to repair all of the DSBs in these experiments, DNA-PKcs and XRCC4 must sequentially localize to breaks, explaining the extended time period during which the chromatin associated fraction of these factors does not change (Fig. S4D). Consistent with this interpretation the duration of the chromatin association plateaus of DNA-PKcs and XRCC4, but not their height, increase with the number of DSBs induced.

### The overall repair capacity of the NHEJ pathway in human cells

To define the rate of NHEJ (i.e. the number of breaks repaired per minute), we used Halo-Ku70 as a marker to determine the number of DNA breaks initially present after exposure to Calicheamicin (Fig. 4D). To measure total repair time, the time from damage induction until dissociation of Halo-XRCC4 was determined, which marks completion of DNA ligation and dissociation of the XRCC4-Ligase 4 complex (Fig. 4B,C). Both the repair time and initial break number increased with increasing concentrations of calicheamicin (Fig. 4B,D). The repair rate after treatment with 20 nM and 40 nM of calicheamicin was identical (~1100 breaks per minute), while the repair rate after break induction with 80 nM was slightly lower (~750 breaks per minute) (Fig. 4E). These observations suggest that the maximal DNA break repair capacity in U2OS cells is approximately 1100 DSBs per minute.

### Analysis of the relative timing of NHEJ factor recruitment reveals kinetics of short-range synaptic complex formation

To precisely determine the relative timing of the recruitment of Halo-Ku70, Halo-DNA-PKcs, Halo-XRCC4, and Halo-XLF to DSBs, time course experiments were performed after induction of DSBs with 40 nM Calicheamicin. While Halo-Ku70 and Halo-DNA-PKcs were recruited to DSBs immediately after break induction, the chromatin recruitment of Halo-XRCC4 and Halo-XLF was delayed by ~6 minutes relative to Halo-Ku70 and Halo-DNA-PKcs (Fig. 5A,B). Halo-XLF was associated with around 600 DNA breaks after DSB induction with 40 nM calicheamicin (Fig. S5A). This suggests that DSBs are initially bound by Ku70/80 and DNA-PKcs and XRCC4 and XLF are recruited at a later time point, consistent with their delayed association observed using LMI (Fig. 3). Halo-DNA-PKcs dissociated from chromatin ~22 minutes after DNA damage induction (Fig. 5A,B). Relative to Halo-DNA-PKcs, Halo-Ku70, Halo-XRCC4, and Halo-XLF remained bound to DSBs for an additional 10 minutes (Fig. 5A,B), consistent with the formation of the ligation competent short-range synaptic complex that lacks DNA-PKcs.

**Figure 5.**
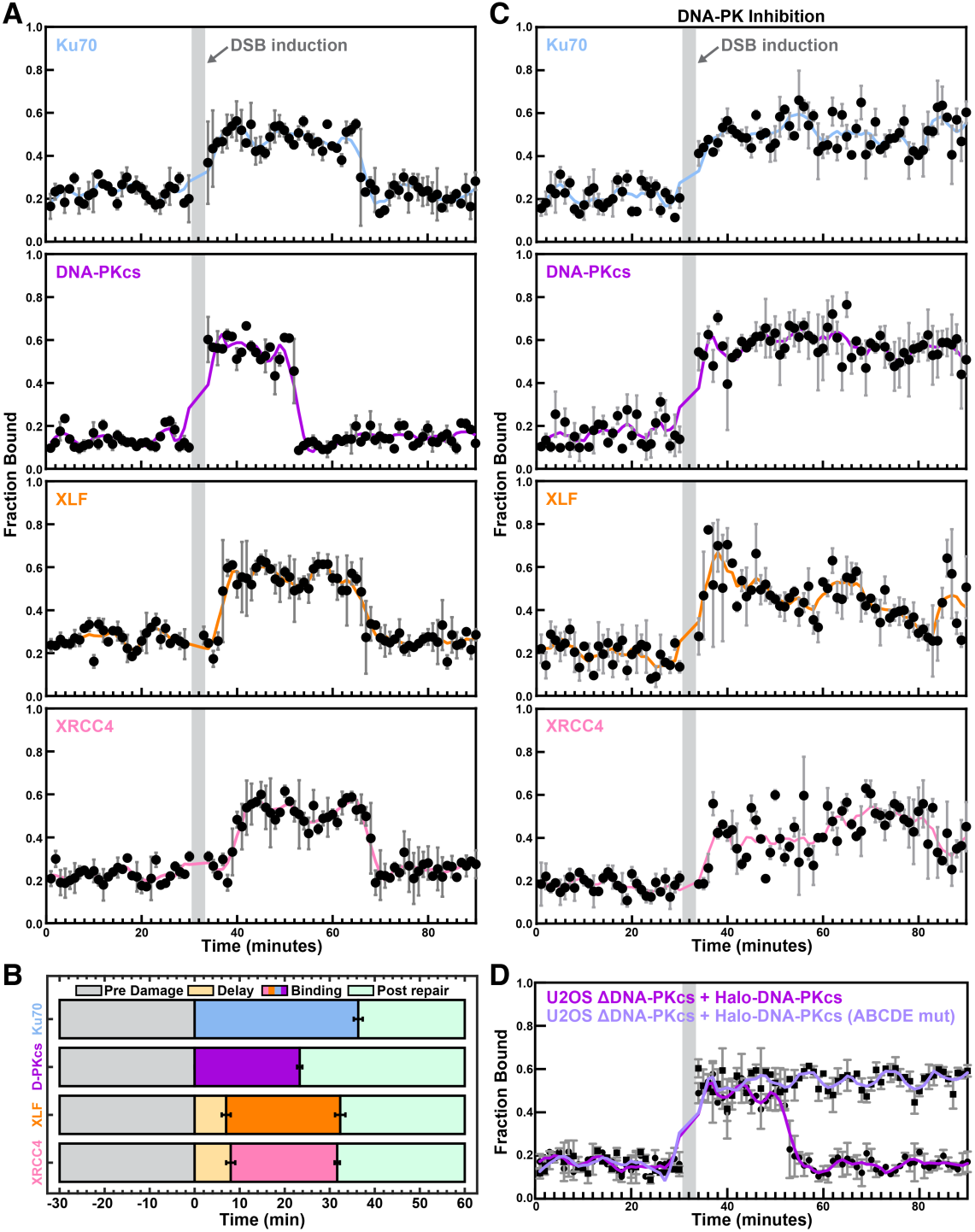
Compositional changes of the NHEJ complex are controlled by DNA-PK activity. **(A)** Plot of the static fraction of each HaloTag NHEJ protein over time. A single cell was imaged at each timepoint. After 30 minutes DSBs were induced with 40 nM calicheamicin for 3 minutes (shaded area) and cells were imaged for 60 minutes thereafter (N = 3, Mean ± S.D.). **(B)** Quantification of the timing of the recruitment of Halo-Ku70, Halo-DNA-PKcs, Halo-XLF, and Halo-XRCC4 after DNA damage induction (N = 3, Mean ± S.D.). **(C)** Plot of the static fraction of each HaloTag NHEJ protein over time in the presence of a DNA-PK inhibitor (Nu7441, 2 nM). A single cell was imaged at each timepoint. After 30 minutes DSBs were induced with 40 nM Calicheamicin for 3 minutes (shaded area) and cells were imaged for 60 minutes thereafter (N = 3, Mean ± S.D.). **(D)** Plot of the static fraction of wildtype Halo-DNA-PKcs or Halo-DNA-PKcs with mutated autophosphorylation sites (ABCDE mut.) expressing in U2OS cells in which DNA-PKcs was knocked out. A single cell was imaged at each timepoint. After 30 minutes DSBs were induced with 40 nM calicheamicin for 3 minutes (shaded area) and cells were imaged for 60 minutes thereafter (N = 3, Mean ± S.D.).

To determine whether the catalytic activity of DNA-PKcs is required for its dissociation from DSBs and in turn the completion of DNA repair marked by the departure of the other NHEJ factors, the experiment was repeated in the presence of an DNA-PK inhibitor (Nu7441) ^37^. When DNA-PKcs kinase activity was inhibited, all four NHEJ factors remained associated with chromatin for >60 minutes (the end of the experiment) (Fig. 5C), suggesting that DNA-PKcs target phosphorylation is required for DNA-PKcs displacement from DSBs (consistent with previous in vitro studies) and potentially short-range complex formation and end ligation. Importantly, this observation is consistent with the requirement for DNA-PKcs catalytic activity for the transition to the short-range complex in single-molecule FRET experiments ^17^. To assess whether DNA-PKcs autophosphorylation at the ADCDE-sites, which is important for DNA-PKcs function ^37–42^, is required for its dissociation from DSBs, wildtype Halo-DNA-PKcs and Halo-DNA-PKcs with the ABCDE sites mutated to alanine (ABCDE mutant) were expressed in DNA-PKcs-deficient U2OS cells. Similar to Halo-DNA-PKcs in the presence of the DNA-PKcs inhibitor, the ABCDE mutant remained associated with chromatin for the entirety of the experiment, while wildtype Halo-DNA-PKcs dissociated from chromatin 20 minutes after DSB induction (Fig. 5D, Fig. S5B). These data are consistent with previous studies showing that ABCDE autophosphorylation is required for DNA-PKcs dissociation from chromatin after DNA damage ^42,43^.

It total, these observations in living cells are consistent with a stepwise maturation of the NHEJ complex at DSBs that requires DNA-PK kinase activity; Ku70/80 and DNA-PKcs initially recognize DSBs and recruit XRCC4, Lig4, and XLF. DNA-PKcs autophosphorylation-induced dissociation occurs well before repair is complete, while the other core NHEJ factors remain associated with the break sites, consistent with progression to the short-range synaptic complexes where final repair occurs.

### Ligase 4 is required for retention of Ku70 and DNA-PKcs at DSBs and its catalytic activity is dispensable for transition to short-range complexes

Ligase 4 has been shown to be necessary for short-range complex formation *in vitro* ^17,44^. To assess the role of Ligase 4 in the maturation of the NHEJ complex in living cells, Ligase 4 was knocked out in cells expressing Halo-tagged Ku70, DNA-PKcs, or XRCC4 and NHEJ factor recruitment to DNA breaks was analyzed after DSB induction with 40 nM Calicheamicin. Re-expression of wild type Ligase 4 recapitulated the recruitment kinetics of Halo-Ku70, Halo-DNA-PKcs, and Halo-XRCC4 observed in the presence of endogenous Ligase 4 (Fig. 6A,B, Fig. 5A, Fig. S5C-E, Movie S11-13). In the absence of Ligase 4, Halo-Ku70 and Halo-DNA-PKcs were recruited to DSBs immediately after induction and simultaneously dissociated from DNA breaks after approximately 16 minutes, substantially faster than in the presence of Ligase 4 (Fig. 6A,B, Fig. 5A, Fig. S5C-E, Movie S11-12). Halo-XRCC4 was not recruited to DSBs in Ligase 4 knockout cells (Fig. 6A, Movie S13), consistent with previous results ^45,46^. These observations demonstrate that Ligase 4 is required for the recruitment of XRCC4 to DNA breaks, and that Ligase 4 stabilizes Ku70 and DNA-PKcs at DSBs, potentially by promoting the stability long-range synaptic complexes (Fig. 1A).

**Figure 6.**
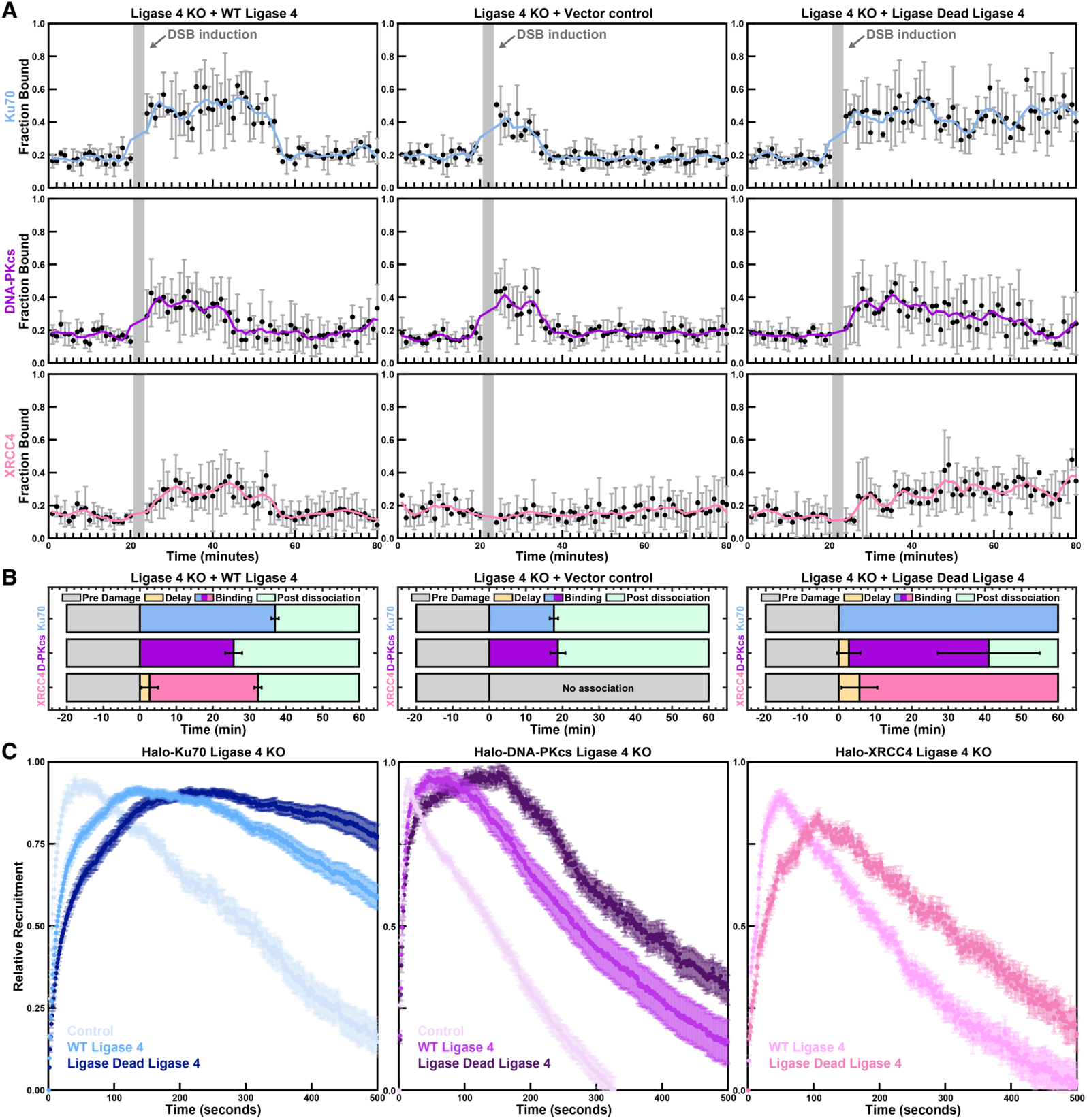
Ligase 4 is required for the retention of Ku70 and DNA-PKcs at DNA breaks. **(A)** Plot of the static fraction of Halo-Ku70 (top row), Halo-DNA-PKcs (middle row), or Halo-XRCC4 (bottom row) over time in ligase 4 knockout cells, transiently expressing wildtype ligase 4 (left column), a vector control (middle column) or ligase dead ligase 4 (right column). After 20 minutes DSBs were induced with 40 nM Calicheamicin for 3 minutes (shaded area) and cells were imaged for 60 minutes thereafter (N = 3, Mean ± S.D.). **(B)** Quantification of the timing of the recruitment of Halo-Ku70, Halo-DNA-PKcs, Halo-XLF, and Halo-XRCC4 after DNA damage induction in Ligase 4 knockout cells expressing a control plasmid, or expression vectors for wildtype and catalytically inactive Ligase 4 (N = 3, Mean ± S.D.). **(C)** Laser micro irradiation of U2OS cells with ligase 4 knockout, expressing Halo-Ku70 (left), Halo-DNA-PKcs (middle), or Halo-XRCC4 (right), transiently expressing wildtype ligase 4, a vector control, or catalytically inactive ligase 4 (N = 30-45 cells per condition, Mean ± S.E.M.).

To dissect the structural and catalytic contributions of Ligase 4 to NHEJ complex formation and maturation, a previously described catalytically inactive allele of Ligase 4 (Ligase 4 5XK) was introduced into Ligase 4 deficient cells (Fig. S5C-E) ^44^. In the presence of catalytically inactive Ligase 4 both Halo-XRCC4 and Halo-Ku70 were retained at DNA breaks for the entire duration of the experiment, demonstrating that DNA ends must be ligated for Ku70 and XRCC4 to be removed from DSBs (Fig. 6A,B). In contrast, DNA-PKcs dissociated from DNA breaks, albeit after a longer delay compared to cell expressing catalytically active Ligase 4 (Fig. 6A, Fig. 5A). This suggests that the short-range synaptic complex can form without the ability to ligate DNA ends and that DNA-PKcs dissociates from breaks prior to the repair event. To confirm these observations, we carried out laser micro-irradiation experiments in Ligase 4 knockout cells expressing Halo-Ku70, Halo-DNA-PKcs, or Halo-XRCC4 transfected with a control vector or expression plasmids encoding wild type Ligase 4 or catalytically inactive Ligase 4 (Movie S14-16). Consistent with the single-molecule imaging experiments, in the absence of Ligase 4, Halo-Ku70 and Halo-DNA-PKcs rapidly dissociated from laser induced DNA lesions, while Halo-XRCC4 was not recruited to sites of DNA damage (Fig. 6C, Fig. S5F-G, Movie S14-16). In addition, Halo-Ku70 was retained for an extended period of time in cells expressing catalytically inactive Ligase 4 (Fig. 6B, Fig. S5F). Importantly, similar to the observation made by single-molecule imaging, Halo-DNA-PKcs dissociated from DNA lesions in the presence of catalytically inactive Ligase 4 (Fig. 6B, Fig. S5F), consistent with the formation of an NHEJ complex that lacks DNA-PKcs when DNA ligation is not possible.

In summary, these observations demonstrate that Ligase 4 is structurally required to retain Ku70 and DNA-PKcs at DSBs and that the short-range synaptic complex that lacks DNA-PKcs can form without the ability to ligate DNA ends.

## DISCUSSION

Non-homologous end joining repairs the vast majority of DNA double strand breaks in human cells. In addition to its central importance in maintaining genome integrity, toxic NHEJ can be unleashed in tumors with defects in other DNA repair pathways to specifically eliminate cancer cells ^1,2^. Therefore, quantitatively defining the molecular mechanisms underlying NHEJ is critical to understand the basic biology of DNA repair and has important clinical implications for targeting DNA repair pathways as a therapeutic approach to treat cancer ^467,48^. Recent single-molecule FRET and Cryo-EM studies suggest that NHEJ is initiated by the formation of a long-range synaptic complex, which can have several distinct compositions that likely exist in equilibrium ^11–15,17^. These long-range complexes transition into short-range synaptic complexes, which facilitate ligation of the DNA ends, in a step that requires DNA-PK kinase activity. While these elegant studies defined critical aspects of the molecular mechanism underlying NHEJ, our knowledge of how NHEJ proceeds in living cells was limited.

Because NHEJ factors are both highly abundant and only small numbers (two to four) of the core NHEJ proteins are recruited to DNA breaks, it has been challenging to detect their recruitment to DSBs in living cells ^20^. In addition, well established methods to analyze the recruitment of DNA repair factors to sites of DNA damage in real time, for example laser micro-irradiation, generate complex DNA lesions, making it difficult to interpret experimental outcomes ^23–25^. To overcome this hurdle, we used single-molecule live-cell imaging to study the kinetics of NHEJ factors recruitment to drug induced DSBs. The key assumption is that the immobilization of individual NHEJ factors in response to treatment with DNA damaging agents is caused by their binding to DSBs. The single-molecule approach has two key advantages: individual molecules can be detected making it possible to visualize the recruitment of factors that are not present at breaks in large numbers, and small molecule drugs can be used to induce a titratable number of chemically defined DNA lesions (as opposed to the complex lesions that result from LMI) ^20, 23–25^,. Using this approach, we analyze the temporal and compositional maturation of NHEJ factors bound to DNA breaks and defined of the overall kinetics and repair capacity of the NHEJ pathway in living cells.

### Repair capacity and overall rate of the non-homologous end joining pathway

Our single molecule imaging-based approach to directly visualize the recruitment of NHEJ factors to DNA breaks allowed enumeration of the chromatin bound DNA repair factors providing an estimate of the overall rate of DSB repair by the NHEJ pathway. A key assumption is that almost all breaks are rapidly bound by Ku70/80. Given the high abundance of Ku70 (> 10^6^ molecules per cell) relative to the estimated number of breaks induced in the experiment (< 10^5^ breaks per cell), this appears to be a reasonable assumption. Using Ku70 binding to DSBs as a read out of the total breaks initially present in cells and the dissociation of the XRCC4-Ligase 4 complex as the time point at which all breaks have been repaired, our experiments indicate that the NHEJ pathway can repair up to 1100 DNA breaks per minute in U2OS cells. Importantly, the repair rate measured after DSB induction with 20 nM and 40 nM of Calicheamicin was identical, with both the break number and repair time doubling. Although the individual measurements in these experiments exhibit variability, the notable concurrence in repair rates across diverse experimental conditions augments our confidence in both the efficacy of our approach and the validity of our interpretation. After damage induction with 80 nM Calicheamicin, a slight reduction in the repair rate to ~750 breaks per minute was observed. While this reduction could simply be a consequence of the large experimental error of this data point it is tempting to speculate that the overall NHEJ repair rate slows down as the number of breaks induced approaches the number of DNA-PKcs molecules present in the cell. Break tethering and long-range synaptic complex formation requires a DNA-PKcs molecule to be bound to each DNA end of a DSB. When cells were exposed to 80 nM Calicheamicin the number of DSBs exceeds 50,000 per cell, which would require 100,000 DNA-PKcs molecules; a number comparable to the abundance of DNA-PKcs determined in this work. Under these conditions it would be possible that repair is delayed because simultaneous binding of DNA-PKcs to both DNA ends is less likely.

While Ku70/80 likely rapidly associates with the vast majority of DNA breaks induced, our results demonstrate that DNA-PKcs only binds to approximately 10000 DSBs, regardless of the number of DNA breaks present. This suggests that a large number of breaks in our experiments are only bound by Ku70/80 or Ku70/80 and DNA-PKcs. In addition, we observed a reduction in the mobility of freely moving DNA-PKcs molecules in the nucleus (D_Free_). This indicates that the properties of DNA-PKcs are changed cell wide, once a substantial number of DNA breaks are present in cells. A key unaddressed question is why DNA-PKcs does not appear to associate with a large fraction of Ku70/80 bound DNA ends. One possibility is that many Ku70/80 bound ends are not ideal substrates for DNA-PKcs because the DNA end is not sufficiently exposed for DNA-PKcs to engage. For instance, if Ku70/80 associates with a DNA end in close proximity to a nucleosome, which might require chromatin remodeling prior to DNA-PKcs binding. However, we currently do not have evidence for this model and future studies are necessary to decipher to mechanism by which DNA-Pkcs is recruited to Ku70/80 bound DNA ends.

In contrast to Ku70 and DNA-PKcs, XRCC4 and XLF were only associated with 1000 and 600 DNA breaks under our experimental conditions, respectively. In combination with the observed repair rate of approximately 1000 breaks per minute per cell, this suggests that the XRCC4-Ligase 4 complex and XLF complete their repair function in 30-60 seconds before moving to the next DNA break. Our results are consistent with the hierarchy of NHEJ factor recruitment to DNA breaks and indicate that the XRCC4-Ligase 4 complex and XLF are the rate limiting factors that control the repair capacity of the NHEJ pathway in human cells. Interestingly, we observed that the majority of Halo-XLF localized to the cytoplasm, rather than the nucleus. In addition, work by others has demonstrated that phosphorylation of XLF by Akt kinase promotes its export to the cytoplasm ^27^. Together, these observations suggest that the regulation of the subcellular localization of XLF could regulate the repair capacity of the NHEJ pathway in human cells.

In total, we conclude that NHEJ has the maximal cellular capacity to repair approximately 1100 DSBs per minute. This rate is controlled by the availability of the XRCC4-Ligase 4 complex and XLF and end repair takes 30-60 seconds once the ligation machinery associates with the DNA break.

### Stepwise maturation of the NHEJ complex in living cells

Our observations demonstrate that Ku70 and DNA-PKcs are recruited to DSBs immediately after their formation. In contrast the recruitment XRCC4 and its constitutive binding partner Ligase 4, along with XLF, is only detectable approximately 6 minutes after the association Ku70 and DNA-PKcs. The observation that Ku70 and DNA-PKcs are targeted before XRCC4 and XLF is consistent with an initial DNA end synapsis by the domain-swap Ku70/80-DNA-PKcs dimer. This dimer can also exist bound to XRCC4, Ligase 4, and XLF, and transitions into the XLF-dependent long-range complex by an unknown mechanism (Fig. 7) ^12–14^. Mutational experiments suggest that short-range complex formation requires prior formation of the XLF-dependent long-range synaptic complex (Fig. 7) ^19^. The four core NHEJ factors remain associated with DSBs for ~15 minutes at which point DNA-PKcs dissociates. We believe that during the 15-minute window in which all NHEJ factors analyzed are observed to be bound to DNA breaks the bulk of end-joining occurs. DNA-PKcs, XRCC4, and XLF transition from one break to the next, explaining why their fraction bound remains constant during this time interval, while the fraction bound of Ku70 gradually decreases as breaks are repaired (Fig. S4D). Once DNA-PKcs dissociates from DNA breaks, Ku70, XRCC4, and XLF remained bound to chromatin for an additional 10 minutes. This suggests that a number of DNA breaks remain in a short-range synaptic complex for an extended period of time, rather than being repaired in 30-60 seconds after XRCC4-Ligase 4 association as outlined above. Since DNA-PKcs binding is no longer observed, it is unlikely that these breaks transition back into a long-range synaptic complex (Fig. 7). We believe these breaks likely require extensive end processing or represent DNA lesions that are incompatible with repair by NHEJ. Once these breaks are repaired or committed to an alternative repair pathway all NHEJ factors dissociate from chromatin. Importantly, the dissociation of DNA-PKcs requires the kinase activity of DNA-PK, specifically DNA-PKcs autophosphorylation at the ABCDE sites, as previously suggested ^41–43^. Moreover, upon inhibition of DNA-PK catalytic activity all core NHEJ factors remain associated with DSBs, demonstrating that short-range synaptic complex formation requires phosphorylation of functionally relevant DNA-PK targets including the ABCDE sites, consistent with the FRET studies ^17^.

**Figure 7.**
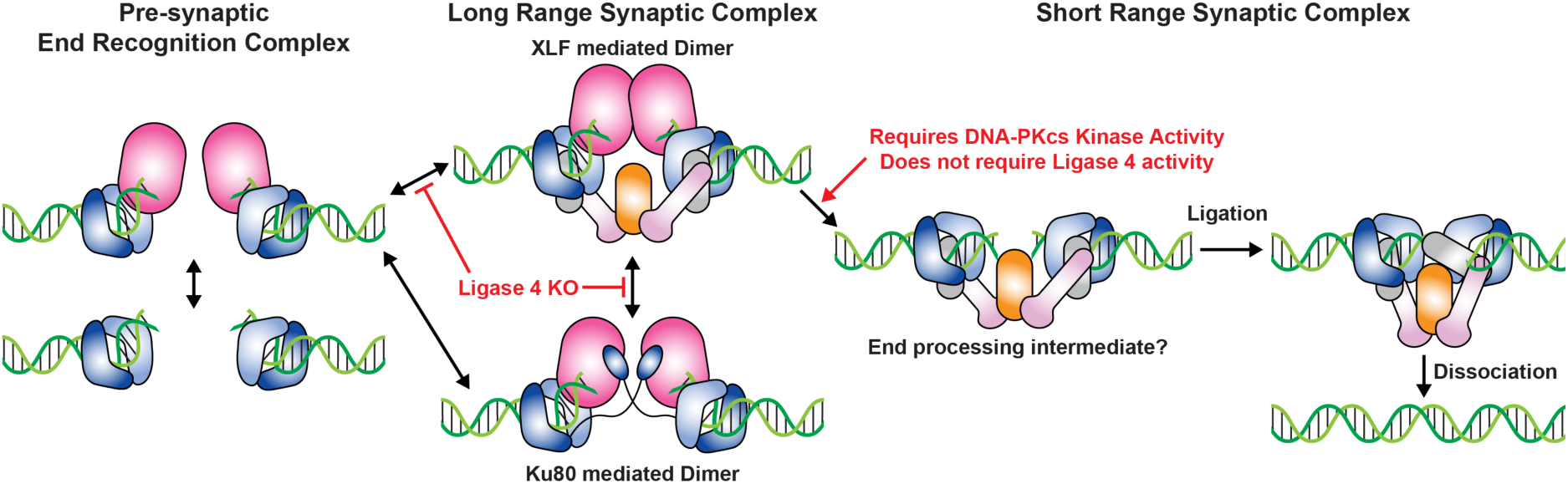
Model for NHEJ. Stepwise maturation of the NHEJ complex throughout DNA break repair, highlighting key transition points controlled by DNA-PKcs catalytic activity and structural contributions of the XRCC4-Ligase 4 complex.

An important limitation of our work is that we cannot determine the time point of long-range complex formation. Graham *et al.* reported that NHEJ long-range complexes are short-lived (only a few seconds) whereas short-range NHEJ complexes are considerably more stable ^17^. We suggest that while DNA-PKcs is associated with DNA ends, numerous long-range complexes form at a single DSB in an iterative fashion to facilitate different aspects of end-processing. Although how each DSB is shuttled between different complexes has not been determined. Emerging data suggest that DNA-PKcs’s unique DNA End Blocking helix, which appears to melt the DNA termini within the DNA-binding cradle, may provide a mechanism for DNA-PK to sample DNA ends so that appropriate NHEJ complexes are formed depending on the end chemistry of the DNA end bound ^19^. Once ligatable, ends are paired within an XLF-mediated dimer, DNA-PKcs autophosphorylates ABCDE sites in trans, resulting in dissociation of DNA-PKcs and progression to the ligation competent short-range complex. Our observations suggest that the transition into this short-range complex that lacks DNA-PKcs does not require the catalytic activity of Ligase 4, consistent with single-molecule FRET studies ^17^. However, DNA-PKcs dissociation is substantially delayed in cells expressing catalytically inactive ligase 4, which suggests that Ligase 4 and potentially its DNA binding activity contributes to the conformational changes required for short-range complex formation, consistent with recent work from Stinson *et al* ^16^.

In total, this study establishes a new methodology to analyze NHEJ factor association with chemically well-defined DNA breaks at the single-molecule level. This approach allowed us to define the overall repair capacity of the NHEJ pathway in human cancer cells and directly visualized the compositional changes in the NHEJ complex during double-strand break repair in living cells.

## ACKNOWLEDGMENTS

We would like to thank members of the Schmidt lab for discussions and Luke Lavis (HHMI Janelia Research Campus) for providing HaloTag dyes. This work was supported by grants from the NIH (DP2-GM142307) to J.C.S., (F32GM139292) to J.R.H., and (R01AI147634) to K.M..

## AUTHOR CONTRIBUTIONS

Conceptualization: M.M, K.M., and J.C.S.; Experiments: M.M., J.R.H., N.J.G., K.M.; Data Analysis: M.M., J.R.H., T. J.; Writing—Original Draft: M.M.; Writing—Review and Editing: M.M., K.M., and J.C.S.

## Materials and Methods

### Cell Lines and Cell Culture

U2OS cells were cultured in RPMI supplemented with 10% fetal bovine serum (FBS), 100 units/mL penicillin, and 100 µg/ml streptomycin. CHO-V3 cells were grown in alpha-MEM containing 10% FBS, 2 mM L-glutamine, 0.1 mM non-essential amino acids, 1 mM sodium pyruvate, 100 U/ml penicillin, 100 µg/ml streptomycin, and 10 µg/ml ciprofloxacin. Stable transfectants of CHO-V3 cell were selected and maintained in complete medium containing 10 μg/ml puromycin. All cells were grown at 37°C in a humidified atmosphere with 5% CO_2_.

### Molecular Cloning, Plasmids and Genome Editing

Endogenous tagging of Ku70, XLF, and XRCC4 was performed with CRISPR/Cas9 mediated homology-directed repair (HDR) to precisely introduce HaloTag. To achieve this, sgRNAs were inserted into the BbsI site of the pX330 plasmid (pX330-U6-Chimeric_BB-CBh-hSpCas9 was a gift from Feng Zhang, Addgene plasmid # 42230) ^49^. Each HDR donor plasmid was was constructed from three DNA fragments by Gibson assembly: a left homology arm (449 bp), a right homology arm (449 bp), and an insertion in-between the homology arms including a 3x FLAG tag, an inverted SV40 promoter, a Puromycin resistance cassette (PuroR) flanked by LoxP sites (for further selection), the HaloTag, and a TEV protease cleavage site, followed by a short peptide linker sequence, as previously described ^50, 51^. Homology arms were ordered as G-blocks from IDT. U2OS cells were transfected with 1 μg of pX330 plasmid and 1 μg of the HDR donor plasmid using 1 ug FuGene 6 (Promega). Approximately 2-3 days post-transfection, edited cells were selected with 1 ug/mL puromycin. These cells were then transfected with 2 μg of plasmid encoding eGFP-Cre recombinase (CAG-GFP-IRES-CRE was a gift from Fred Gage, Addgene plasmid # 48201) ^52^. To obtain individual clones cells were labeled with JFX650, and single cells that were eGFP and JFX650 positive were sorted into 96-well cell culture plates containing 50% conditioned media (media supernatant from previous cultures of U2OS cells filtered through a 0.2 µm filter). For knockout of DNA-PKcs, cells were transfected with two gRNA plasmids (TGCAACTTCACTAAGTCCA, GAAAAAGTACATTGAAATT). Knockout was confirmed by Western blot and Sanger sequencing. For transient transfection of wild type Halo-DNA-PKcs and Halo-ABCDE plasmids, 1ug of plasmid and FuGene 6 (Promega) were used. CHO-V3 transfectants expressing human DNA-PKcs were derived by stably transfecting cells with 10 ug expression plasmid and 1 ug pSuper-Puro with PEI. 48 hours after transfection, cells were plated at limiting dilution in media containing 5ug/ml puromycin. Isolated clones were screened for DNA-PKcs expression by immunoblotting. Ligase 4 KO cell lines were generated using a gRNA and Cas9 nuclease targeting the coding region of Ligase 4. The gRNA (CATACGTTCACCATCTAGCT) was previously cloned into pCas2a-Puro. 500 ng of plasmid was transfected into 5*10^5^ cells using a Neon nucleofector (ThermoFisher, 4×10 ms pulses at 1230 V) and given 48 hours to recover. Clones were selected by plating limiting dilutions into 10 cm dishes in complete media + 2 ug/mL puromycin for 48 hours. After colonies had grown to ~50 cell size, individual clones were manually transferred to a 24-well plate and allowed to grow to confluency. Ligase 4 KO clones were analyzed by western blot probed with anti-Ligase 4 (Abcam, 26039). Ligase 4 was transiently complemented into KO cell lines by transfecting 500,000 cells with 100 ng of pmaxGFP (Lonza) and 500 ng of expression plasmid (empty vector, pcDNA3-Ligase 4 WT, or pcDNA3 catalytically inactive Ligase 4 described as 5xK-R in Goff et al. 2022) ^44^. Approximately 100,000 cells/well were plated into 24-well glass-bottom dishes (Cellvis, P24-1.5H-N) and allowed to recover for 48-72 hours before imaging.

### Antibodies and immunoblotting

Cell extracts were prepared in 2x Laemmli buffer with β-mercaptoethanol. Proteins were resolved by SDS–PAGE using Mini-PROTEAN TGX stain-free gels (BioRad), except for DNA-PKcs for which homemade 6% polyacrylamide gels were used. Proteins were transferred to PVDF using either a Trans-Blot Turbo system (with Turbo transfer buffer) (BioRad) or by traditional wet tank transfer using CAPS Buffer with 10% Methanol (for DNA-PKcs). Antibodies used in this study were anti-DNA-PKcs (1:1000 dilution) ^53^, anti-XLF (Invitrogen PA596310, 1:1000), anti-XRCC4 (Invitrogen, PA5-82264, 1:2000), anti-Ku70 (Invitrogen, MA5-12933, 1:1000), anti-Lamin B1 (Proteintech, 12987-1-AP), and anti-Tubulin (Proteintech, 66031-1-Ig). HRP-conjugated secondary antibodies were used at 1:5000 dilutions and chemiluminescence signal was produced using the Clarity western ECL substrate (BioRad) and detected using a ChemiDoc imaging system (BioRad). All western blots were carried out in at least 3 technical replicates and were analyzed using ImageQuant TL 8.2.

### Measurement and quantification of in-gel fluorescence and protein abundance

To measure total protein abundance by in-gel fluorescence, cells were labeled for 30 minutes with 500 nM JF646, JFX650, or JF657 HaloTag ligand. For the analysis the labeling efficiency during single-molecule imaging experiments cells were either labeled for 30 minutes with 500 nM JFX650 to determine the total number of HaloTagged NHEJ factors present or using single-molecule imaging labeling conditions (Halo-Ku70 100 pM for 30 seconds, Halo-DNA-PKcs 1 nM for 1 minute, Halo-XRCC4 50 nM for 1 minute, Halo-XLF 10 nM for 1 min). Samples were loaded and separated on Mini-PROTEAN TGX stain-free gels (BioRad) and fluorescence and then total protein were detected using a BioRad Chemidoc, using the Cy5.5 filter and stain-free gel detection after 45 seconds of UV activation, respectivly. For in-gel fluorescence quantification of the HaloTagged proteins, we used a previously described method ^26,54^. Briefly, we created a standard curve using known amounts of recombinant 3X FLAG-HaloTag labeled with JF646, or JF657 and cell lysates from a known number of U2OS cells to evaluate protein abundance in two clones for each cell line. To account for the possibility of different expression of HaloTagged proteins relative to untagged proteins, we determined the ratio of HaloTagged relative to endogenous protein expression using western blot. All in-gel fluorescent SDS-page gels were carried out in at least 3 technical and 3 biological replicates. All gels were analyzed using ImageQuant TL 8.2. For quantification of the labeling efficiency in single-molecule imaging experiments, the fluorescence signal was normalized to the total protein signal to normalize for differences in loading and cell number.

### Clonogenic Survival Assays

The day before zeocin treatment, 500 cells were seeded in six-well plates in triplicate. The next day, cells were treated using a range zeocin (Gibco) concentrations for four hours, and then media with zeoncin was replaced with fresh complete media without drug. Approximately 7–10 days after exposure to zeocin, 6-well plates were rinsed with PBS, and cells were fixed and stained in crystal violet solution (20% ethanol and 1% w/v crystal violet). The crystal violet solution was removed by carefully careful washing with water. Dry plates were imaged using the Coomassie Blue filter set on a BioRad Chemidoc. Colony counts were determined using ImageQuant TL 8.2. All colony survival assays were performed with 4 biological replicates.

### Live cell imaging

Live cell imaging was carried out on a Olympus IX83 inverted microscope equipped with a 4-line cellTIRF illuminator (405 nm, 488 nm, 561 nm, 640 nm lasers), an Excelitas X-Cite TURBO LED light source, an Olympus UAPO 100x TIRF objective (1.49 NA), an Olympus UPlanApo 60x TIRF oil-immersion objective (1.50 NA), a CAIRN TwinCam beamsplitter, two Andor iXon Ultra EMCCD cameras or two Hamamatsu Orca BT Fusion cameras, a cellFRAP with a 100 mW 405 nm laser, an environmental control enclosure for temperature, CO_2_, and humidity control. The microscope was operated using the Olympus cellSense software.

### Laser Micro-irradiation

Cells were labeled with the Halo-tag ligand JFX650 (a kind gift from the Lavis lab, Janelia Farms, Ashburn, VA) at a concentration of 500 nM in complete media for 30 min at 37°C, washed three times, and pre-sensitized with Hoechst (1 µg/ml) for ten minutes. The 405-nm laser was directly coupled to the epifluorescence optical path of the microscope with 100x or 60x objective and images were acquired every second using the 630 nm LED light source. DNA lesions were generated in a straight line across the nucleus by micro-irradiation with the 405-nm laser at 25% laser power using 20 ms pulse. For quantitative analyses, we first cropped cells and drift corrected using NanoJ in Fiji ^55^. Intensity profiles of the laser micro-irradiated stripe were generated using ImageJ and data was analyzed using GraphPad Prism by fitting an exponential one-phase association model and one-phase decay model. For association phase we fit the data points from damage induction until maximal binding was observed. For dissociation we only considered data from the time point of maximal binding until the end of the experiment.

### Single-molecule imaging

Cells were grown on 24-well glass bottom dishes (Cellvis, P24-1.5H-N) for 24 hours prior to imaging. Cells were sparsely labeled with JFX650 or JF646 dye ^56,57^ in complete media (Halo-Ku70 100 pM for 30 seconds, Halo-DNA-PKcs 1 nM for 1 minute, Halo-XRCC4 50 nM for 1 minute, Halo-XLF 10 nM for 1 min) and were washed three times with fresh media and incubated for an additional 15 min at 37°C (5% CO_2_) to remove excess dye. Cells were treated with 100 ug/mL zeocin or DMSO for one hour prior to HaloTag labeling and imaged at 146 frames per second for 3000 frames using the Andor 897 Ultra EM-CCD camera. For calicheamicin treatment, cells were labeled and imaged for 30 minutes prior to addition of calicheamicin for 3 minutes while in cellVivo chamber. Calcicheamicin was removed with at least 3 quick washes with complete media. To inhibit DNA-PK cells were treated with 100 nM Nu7441 for 2 hours prior to and during imaging. Cells for imaged at 179 frames per second for 1500 frames using the Hamamatsu Orca Fusion BT sCMOS camera. All experiments were carried out in triplicate or more with at least 15 cells analyzed for each experiment.

### Analysis of Live-Cell Imaging

To analyze the single-molecule particles TIFF files, we used the batch parallel-processing version of SLIMFAST in MatLab 2020b ^32^. For tracking, the following settings were used for all the proteins: Exposure Time = 5.6 or 6.8 ms, NA = 1.49 or 1.50, PixelSize = 0.16 or 0.1083 µm, Emission Wavelength = 664 nm, D_max_ = 5 µm^2^ s^−1^, Number of gaps allowed = 2, Localization Error = 10^−5^, Deflation loops = 0. We only analyzed particles located in nucleus using bright field images to make a mask of nucleus in ImageJ. Using tracked particles, we determined the diffusion coefficients and the bound fractions using the MATLAB version of SpotOn ^32^. The following settings were used: TimeGap = 5.6 or 6.8 ms, dZ = 0.700 µm, GapsAllowed = 2, TimePoints = 8, JumpsToConsider = 4, BinWidth = 0.01 µm, PDF fitting. We applied a 2-state model with the following settings: D_Free2_1State = [0.5 5], D_Bound2_State = [0.0001 0.5] with an assumption that NHEJ factors are or chromatin bound or are freely diffusing. To compare diffusion coefficients and bound fractions, we performed by two-way ANOVA analysis with Tukey’s posthoc test in GraphPad Prism. We analyzed either trajectories from individual cells or pooled sets of cells as indicated in the text.

### Quantification of DNA break number and NHEJ rate

The number of DNA breaks bound be each DNA repair factor analyzed by single-molecule imaging was calculated by determining the increase in the fraction bound after DNA damage induction (F_bound pre damage_ – F_bound post damage_). The F_bound pre damage_ was determined by pooled SpotOn analysis of all cells imaged prior to damage. The F_bound post damage_ was determined by pooled SpotOn analysis of the first 5 cells post damage for Halo-Ku70, or the first 10 cells after chromatin recruitment was observed for Halo-DNA-PKcs, Halo-XRCC4, and Halo-XLF. We only used 5 cells for Halo-Ku70 because the F_bound_ for Halo-Ku70 rapidly decreased after initial recruitment, while it remained constant for extended periods of time for the other factors analyzed. The fraction bound was then multiplied by the average number of single molecule localizations per imaging frame to determine the number of bound molecules observed, which was also provided by the pooled SpotOn analysis. The number of bound molecules was then divided by the labeling efficiency for each protein (Fig. S3B). Because the majority of the nucleus is outside of the focal plane of the objective used in this experiment, we further adjusted our number to account for the subsection of the nucleus images. The central 1 µm thick slice of a sphere with a 4 µm diameter is 1/2.7 of the total volume of the sphere. Since we cannot be certain that we always imaged the center of the nuclear volume we divided by 1/3 rather than 1/2.7 to account for the reduction in the fraction of the nucleus imaged when not imaging the exact center of the nucleus. Finally, the number was divided by the number of repair factors bound to each DNA break.

### Quantification of the timing of NHEJ factor recruitment to DNA breaks

Bound fractions from single-molecule analysis were analyzed using custom MATLAB script. The algorithm preprocessed the fraction bound values using moving average filter. Next, the threshold was determined as a mean plus standard deviation of fraction bound values from all replicas before the drug treatment. Finally, the start and end point of DNA repair state were detected using sliding window with a length of 3 if the median of values within the window was above or below the threshold, respectively.

### Cellular fractionation

Cellular fractionation was performed as previously described ^58^. Briefly, 20 million cells were trypsinized, collected, and washed in ice-cold PBS before swelling in a hypotonic buffer (10 mM HEPES pH 7.9, 10 mM KCl, 1.5 mM MgCl_2_, and 0.5 mM DTT). Then, using a pre-cooled dounce homogenizer and the tight pestle, swollen cells were ruptured with 13 strokes (VWR Cat. 62400-595). The purify the nuclei, burst cells were centrifuged at 218xg for 5 min. To further improve the quality of purified nuclei, the nuclear pellet was resuspended in buffer S1 (0.25 M sucrose, 10 mM MgCl_2_), layered on top of buffer S2 (0.35 M sucrose, 0.5 mM MgCl_2_) in a 15 ml falcon tube, and centrifuged at 1430xg in a swinging bucket rotor for 5 min. Nuclei then were resuspended in buffer S2. Samples of each fraction were analyzed by western blot.

## SUPPLEMENTAL MATERIAL

**Supplemental Figure 1.**
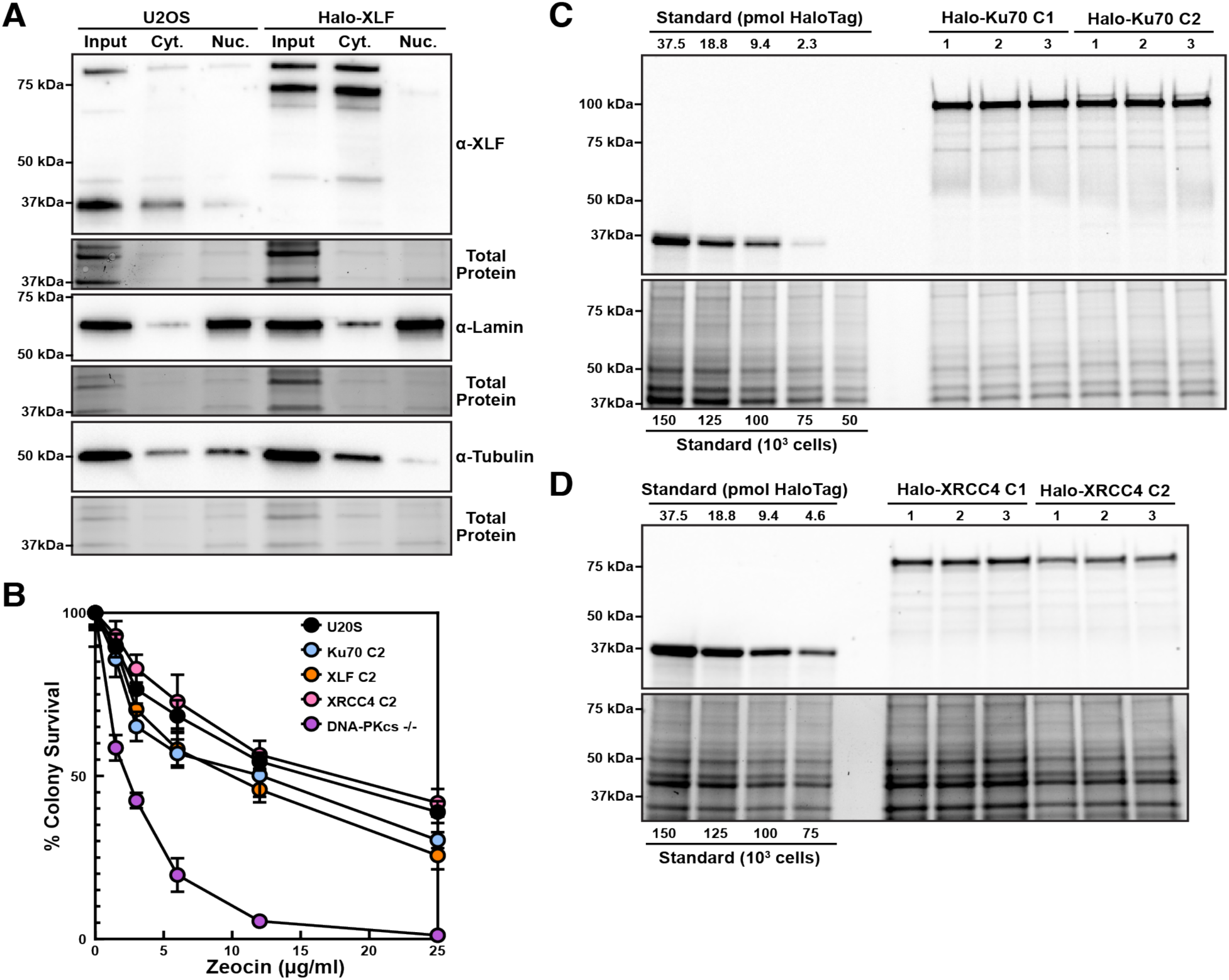
Analysis of sub-cellular localization, functionality, and total abundance of HaloTagged DNA repair factors. **(A)** Western blot analysis of cytoplasmic (Cyt.) and nuclear (Nuc.) fractions from parental U20S and Halo-XLF cells probed with antibodies against XLF, Lamin (nuclear marker), and Tubulin (cytoplasmic marker). **(B)** Clonogenic survival assay of of a second independent clone of for all U2OS cells expressing HaloTagged NHEJ factors, parental U2OS cells, and U2OS cells with DNA-PKcs knockout after challenge with zeocin (N = 4, 3 technical replicates per biological replicate, Mean ± S.D.). **(C-D)** Fluorescence imaging (JF646) and total protein staining for two clones of **(C)** Halo-Ku70 and **(D)** Halo-XRCC4 expressing cells, alongside a protein standard containing a known amount of recombinant 3xFLAG-HaloTag protein labeled with JF646 and whole cell lysate of a known number of cells.

**Supplemental Figure 2.**
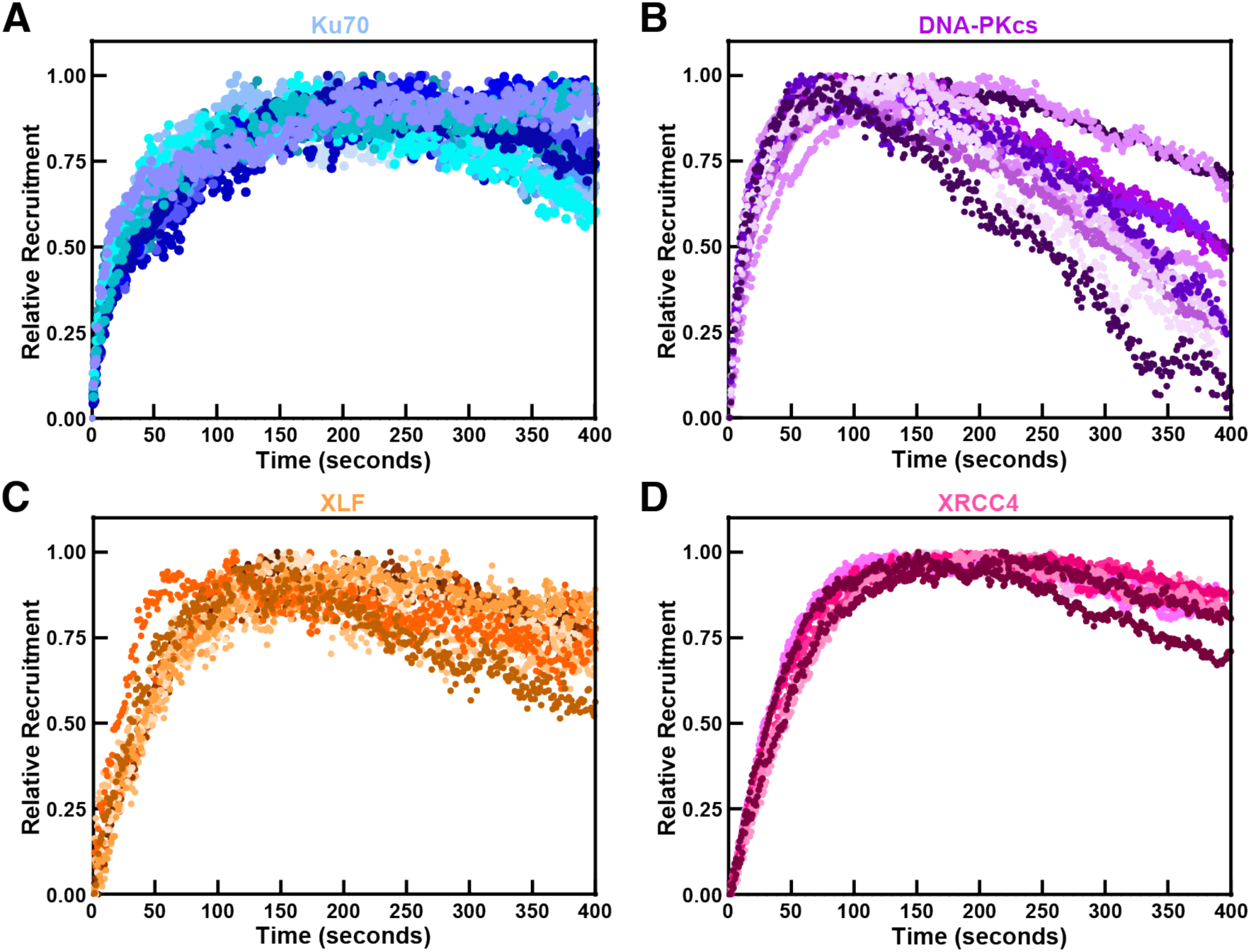
Analysis of HaloTagged NHEJ factor recruitment to LMI induced DNA lesions. **(A-D)** Plots showing normalized fluorescence intensity and recruitment of **(A)** Halo-Ku70, **(B)** Halo-DNA-PKcs, **(C)** Halo-XLF, and **(D)** Halo-XRCC4 to laser-induced DNA lesions. Each shade represents data from an individual cell that was analyzed, and the fluorescence intensity was normalized to the brightest frame of each accumulation curve.

**Supplemental Figure 3.**
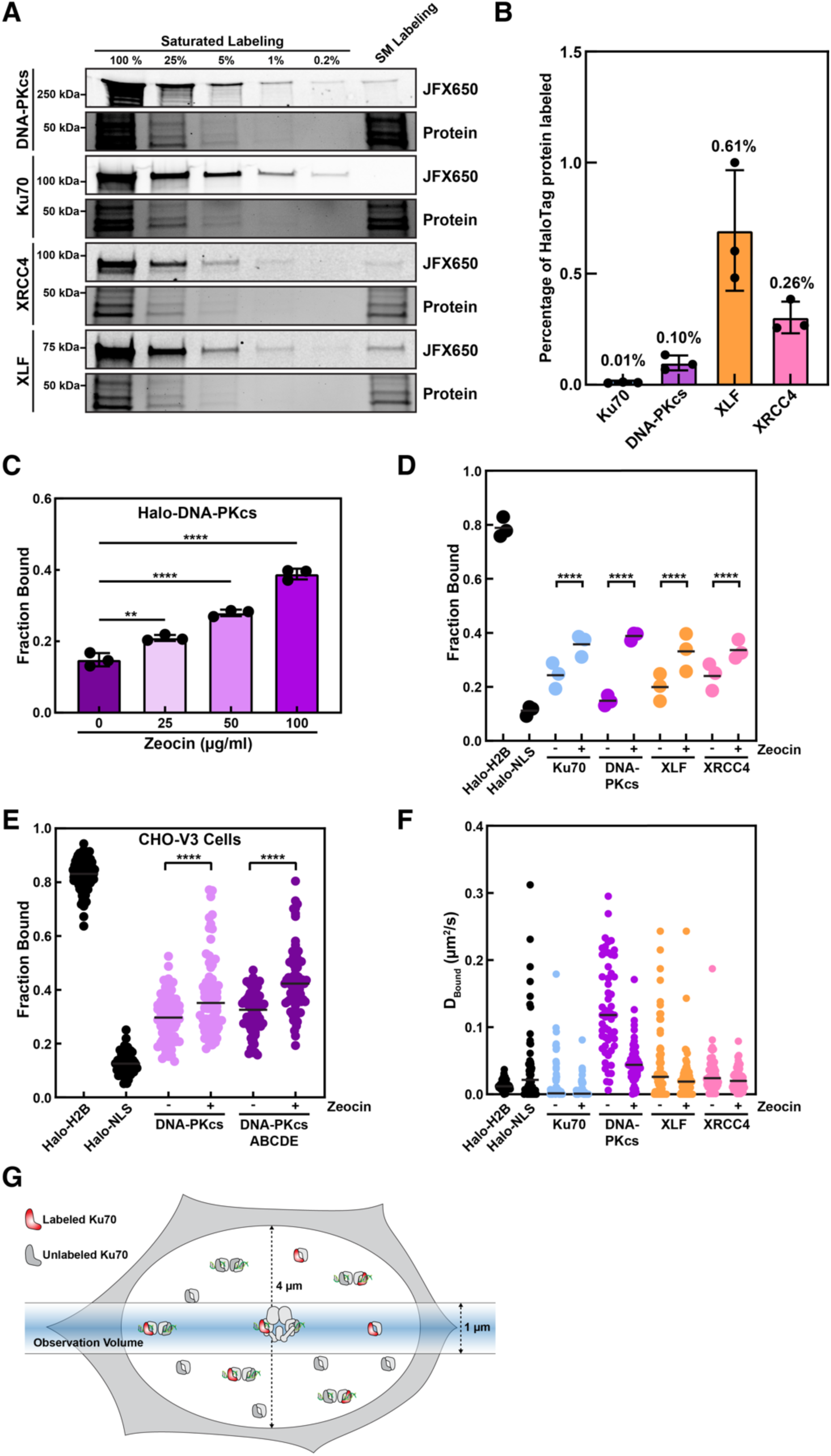
Labeling analysis and single-particle tracking of HaloTagged NHEJ factors. **(A)** Fluorescence imaging and total protein staining of an SDS-PAGE gel with a standard of each HaloTagged NHEJ factor after labeling under saturation conditions (30 minutes, 100 nM HaloTag ligand) or partial labeling for single-molecule imaging (30 seconds for Halo-Ku70, 1 minute for Halo-DNA-PKcs, Halo-XRCC4, and Halo-XLF, 10 nM HaloTag ligand) with JFX650 HaloTag ligand. **(B)** Quantification of the fluorescence intensity normalized to total protein loading for the gels shown in **(A)** (Mean ± S.D.). **(C)** Fraction bound of Halo-DNA-PKcs in the presence of increasing amounts of Zeocin. Each data point represents the Spot-On fit of the pooled data from all cells in a biological replicate (N = 3, n ≥ 20 cells for each protein per replicate and condition, Black bar = median). Data were analyzed by two-way ANOVA with Tukey’s posthoc test (** = p < 0.01, **** = p < 0.0001). **(D)** Plot of the Fraction Bound for each HaloTag NHEJ protein under untreated conditions and post-Zeocin exposure that were analyzed using a two-state model of diffusion. Each data point represents pooled data from all cells in a biological replicate. (N = 3, n ≥ 20 cells for each protein per replicate and condition, Black bar = median). Data were analyzed by two-way ANOVA with Tukey’s posthoc test (**** = p < 0.0001). Same data presented in Figure 3B but analyzed by Spot-On using the pooled step size distribution from all cells for each biological replicate. **(E)** Plot of the Fraction Bound for Halo-DNA-PKcs wild type and Halo-DNA-PKcs ABCDE mutant transiently expressed in CHO-V3 cells under untreated conditions and post-Zeocin exposure that were analyzed using a two-state model of diffusion. Each dot represents the fraction bound of each protein in an individual cell (N = 3, n ≥ 20 cells for each protein per replicate and condition, Black bar = median). Data were analyzed by two-way ANOVA with Tukey’s posthoc test (**** = p < 0.0001). **(F)** Diffusion coefficients of static molecules of HaloTag DDR proteins. Each data point represents the D_Bound_ calculated from the tracks in an individual cell (N = 3, n ≥ 20 cells for each protein per replicate and condition, Black bar = median). **(G)** Graphical rational for the calculation of the total number of breaks bound by the DNA repair factors in the single-molecule imaging experiments.

**Supplemental Figure 4.**
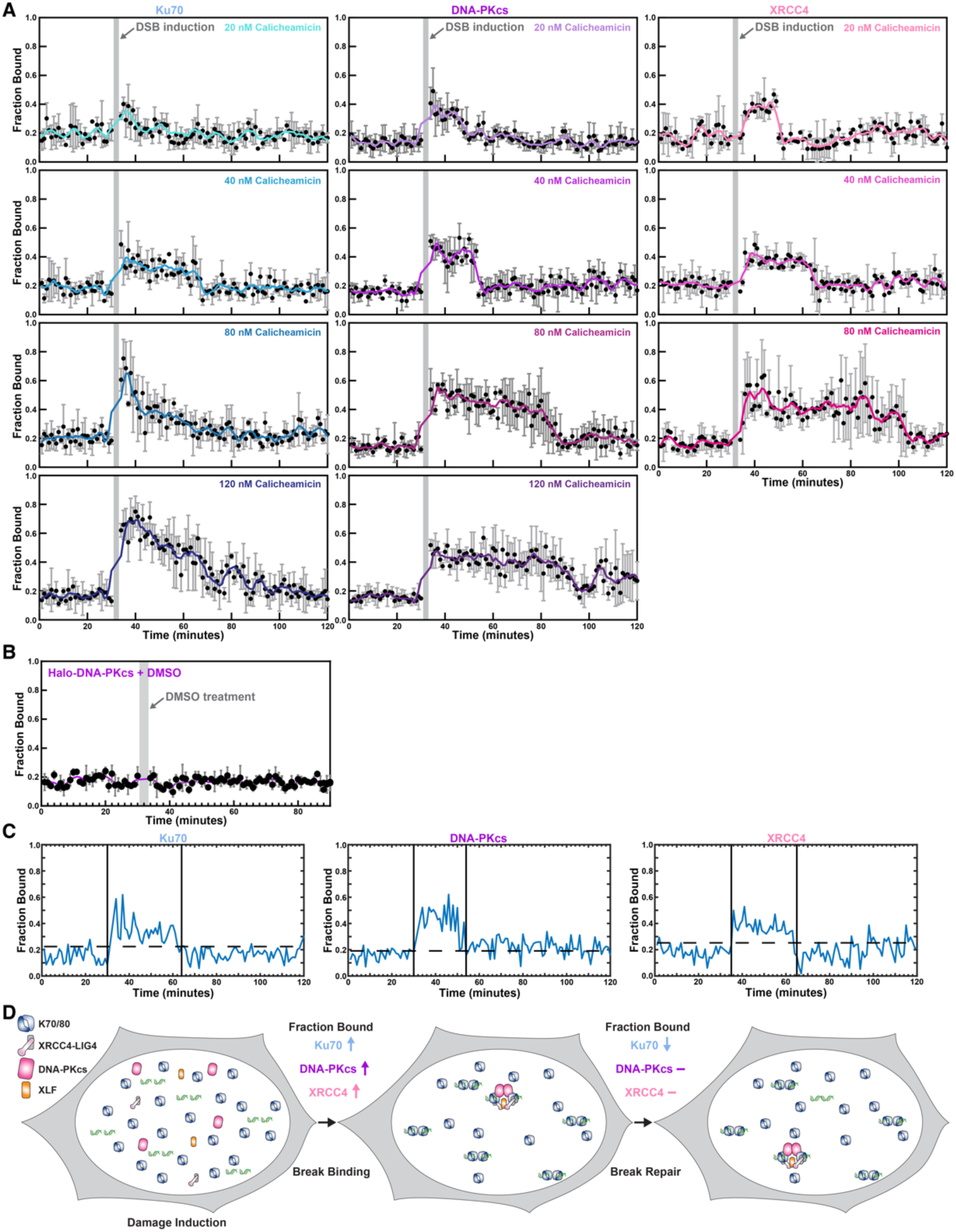
Labeling analysis and single-particle tracking of HaloTagged NHEJ factors. **(A)** Plot of the static fraction of Halo-Ku70, Halo-DNA-PKcs, and Halo-XRCC4 before and after DNA damage induction (shaded area) with various concentrations of Calicheamicin for 3 minutes (N = 3, Mean ± S.D). Lines represent the rolling average over three consecutive timepoints. **(B)** Plot of the static fraction of Halo-DNA-PKcs after treatment with DMSO for 3 minutes (shaded area, N = 3, Mean ± S.D). **(C)** Quantification of the time window of chromatin association for Halo-Ku70, Halo-DNA-PKcs, and Halo-XRCC4. The time between the vertical bars is considered the binding time, the horizontal dashed bar indicates the binding threshold. **(D)** Model for the consecutive repair of DNA breaks in our time course experiments. Because Ku70 is present in large excess relative to DNA-PKcs and XRCC4 its binding is expected to be reduced when breaks are repaired, while the static fraction of DNA-PKcs and XRCC4 remains constant, when the number of breaks present exceed the number of DNA-PKcs and XRCC4 molecules present in the cell.

**Supplemental Figure 5.**
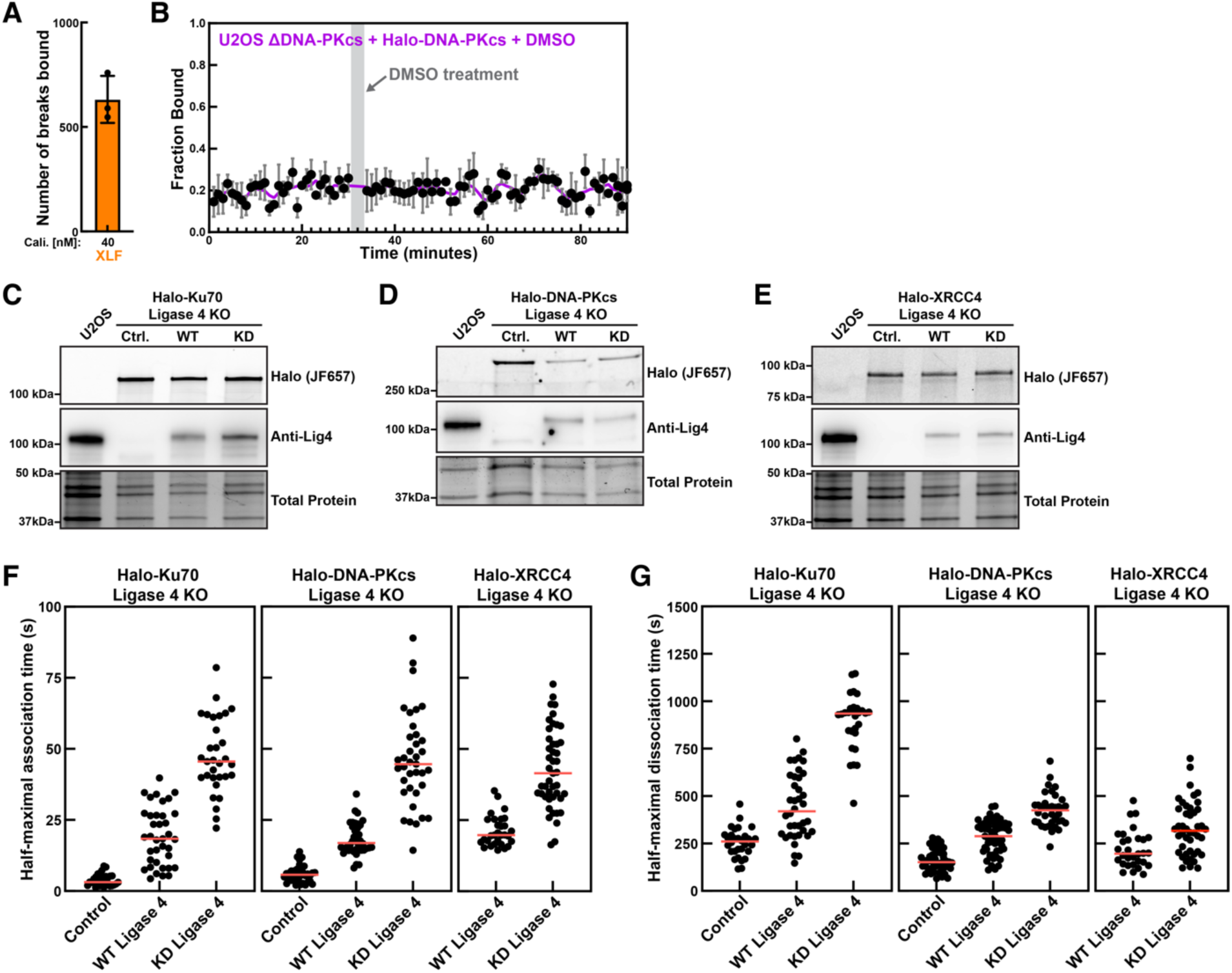
Kinetics of chromatin recruitment of Halo-DNA-PKcs under control conditions. **(A)** Quantification of the number of break bound Halo-XLF (pooled data for 10 time points after chromatin recruitment) molecules after DNA damage induction with 40 nM calicheamicin (N = 3, Mean ± S.D.). **(B)** Plot of the static fraction of Halo-DNA-PKcs in U2OS cells with endogenous DNA-PKcs knocked out and Halo-DNA-PKcs expressed by stable transfection after treatment with DMSO (shaded area, N = 3, Mean ± S.D.). **(C-E)** Fluorescence gels (top), western blots (middle), and total protein stain of cell lysates generated from parental U2OS cells, and U2OS cells expressing **(C)** Halo-Ku70, **(D)** Halo-DNA-PKcs, or **(E)** Halo-XRCC4 labeled with JF657 Halo-ligand in which endogenous ligase 4 was knocked out, transiently transfected with a control vector (Ctrl.) of expression vectors for wildtype ligase 4 (WT) or catalytically inactive ligase 4 (KD). **(F-G)** Half-maximal association **(F)** and dissociation times **(G)** for laser micro irradiation experiments of U2OS cells with ligase 4 knockout, expressing Halo-Ku70 (left), Halo-DNA-PKcs (middle), or Halo-XRCC4 (right), transiently expressing wildtype ligase 4, a vector control, or ligase dead ligase 4 (N = 30-45 cells per condition, Mean). Each data point represents the fit of the data derived from an individual cell. Half-maximal association and dissociation times were fit by using only the data prior or after maximal fluorescence accumulation for each cell, respectively.

## SUPPLEMENTAL MOVIE LEGENDS

**Movie S1.** Representative movie demonstrating recruitment of JFX650-labeled 3xFLAG-HaloTagged Ku70 to DNA DSBs after laser microirradiation. Images were acquired at one frame per second.

**Movie S2.** Representative movie demonstrating recruitment of JFX650-labeled 3xFLAG-HaloTagged-DNA-PKcs to DNA DSBs after laser microirradiation. Images were acquired at one frame per second.

**Movie S3.** Representative movie demonstrating recruitment of JFX650-labeled 3xFLAG-HaloTagged XRCC4 to DNA DSBs after laser microirradiation. Images were acquired at one frame per second.

**Movie S4.** Representative movie demonstrating recruitment of JFX650-labeled 3xFLAG-HaloTagged XLF to DNA DSBs after laser microirradiation. Images were acquired at one frame per second.

**Movie S5.** Representative live-cell single-molecule imaging movie of Halo-NLS transiently expressed in U2OS cells, labeled with JFX650, and acquired at 179 frames per second using the Hamamatsu ORCA BT Fusion camera.

**Movie S6.** Representative live-cell single-molecule imaging movie of Halo-H2B transiently expressed in U2OS cells, labeled with JFX650, and acquired at 179 frames per second using the Hamamatsu ORCA BT Fusion camera.

**Movie S7.** Representative live-cell single-molecule imaging movie of 3xFLAG-HaloTagged Ku70 in a control cell (left) and a cell treated with zeocin (right), labeled with JFX650, and acquired at 146 frames per second using the Andor 897 Ultra camera.

**Movie S8.** Representative live-cell single-molecule imaging movie of 3xFLAG-HaloTagged XLF in a control cell (left) and a cell treated with zeocin (right), labeled with JFX650, and acquired at 146 frames per second using the Andor 897 Ultra camera.

**Movie S9.** Representative live-cell single-molecule imaging movie of 3xFLAG-HaloTagged XRCC4 in a control cell (left) and a cell treated with zeocin (right), labeled with JFX650, and acquired at 146 frames per second using the Andor 897 Ultra camera.

**Movie S10.** Representative live-cell single-molecule imaging movies of 3xFLAG-HaloTagged DNA-PKcs labeled with JFX650 before and after treatment with calicheamicin, acquired at 179 frames per second using the Hamamatsu ORCA Fusion BT camera.

**Movie S11.** Representative single-molecule imaging movie of Halo-Ku70 in Ligase 4 knockout out cells expressing wild-type Ligase 4 (top row), a vector control (middle row), or catalytically inactive Ligase 4 (bottom row) before DNA damage induction with calicheamicin (left column), immediately after DNA damage induction (middle column), and 35 minutes after DNA damage induction (right column) imaged at 171 frames per second using the Hamamatsu ORCA Fusion BT camera.

**Movie S12.** Representative single-molecule imaging movie of Halo-DNA-PKcs in Ligase 4 knockout out cells expressing wild-type Ligase 4 (top row), a vector control (middle row), or catalytically inactive Ligase 4 (bottom row) before DNA damage induction with calicheamicin (left column), immediately after DNA damage induction (middle column), and 35 minutes after DNA damage induction (right column) imaged at 171 frames per second using the Hamamatsu ORCA Fusion BT camera.

**Movie S13.** Representative single-molecule imaging movie of Halo-XRCC4 in Ligase 4 knockout out cells expressing wild-type Ligase 4 (top row), a vector control (middle row), or catalytically inactive Ligase 4 (bottom row) before DNA damage induction with calicheamicin (left column), immediately after DNA damage induction (middle column), and 35 minutes after DNA damage induction (right column) imaged at 171 frames per second using the Hamamatsu ORCA Fusion BT camera.

**Movie S14.** Representative laser micro-irradiation movie of Halo-Ku70 in Ligase 4 knockout out cells expressing wild-type Ligase 4 (left panel), a vector control (middle panel), or catalytically inactive Ligase 4 (right panel) imaged at 1 frame per second using the Hamamatsu ORCA Fusion BT camera.

**Movie S15.** Representative laser micro-irradiation movie of Halo-DNA-PKcs in Ligase 4 knockout out cells expressing wild-type Ligase 4 (left panel), a vector control (middle panel), or catalytically inactive Ligase 4 (right panel) imaged at 1 frame per second using the Hamamatsu ORCA Fusion BT camera.

**Movie S16.** Representative laser micro-irradiation movie of Halo-XRCC4 in Ligase 4 knockout out cells expressing wild-type Ligase 4 (left panel), a vector control (middle panel), or catalytically inactive Ligase 4 (right panel) imaged at 1 frame per second using the Hamamatsu ORCA Fusion BT camera.5

## REFERENCES

1. Chapman, J.R., Taylor, M.R.G., and Boulton, S.J. (2012). Playing the End Game: DNA Double-Strand Break Repair Pathway Choice. Mol Cell 47. 10.1016/j.molcel.2012.07.029.

2. Stinson, B.M., and Loparo, J.J. (2021). Repair of DNA Double-Strand Breaks by the Nonhomologous End Joining Pathway. Annu Rev Biochem 90. 10.1146/annurev-biochem-080320-110356.

3. Watanabe, G., and Lieber, M.R. (2023). The flexible and iterative steps within the NHEJ pathway. Prog Biophys Mol Biol 180–181, 105–119. 10.1016/j.pbiomolbio.2023.05.001.

4. Karanam, K., Kafri, R., Loewer, A., and Lahav, G. (2012). Quantitative Live Cell Imaging Reveals a Gradual Shift between DNA Repair Mechanisms and a Maximal Use of HR in Mid S Phase. Mol Cell 47. 10.1016/j.molcel.2012.05.052.

5. Jasin, M., and Rothstein, R. (2013). Repair of strand breaks by homologous recombination. Cold Spring Harb Perspect Biol 5. 10.1101/cshperspect.a012740.

6. Torgovnick, A., & Schumacher, B. (2015). DNA repair mechanisms in cancer development and therapy. In Frontiers in Genetics (Vol. 6, Issue APR). 10.3389/fgene.2015.00157

7. Loparo, J. J. (2023). Holding it together: DNA end synapsis during non-homologous end joining. DNA Repair, 130. 10.1016/j.dnarep.2023.103553

8. Grundy, G.J., Moulding, H.A., Caldecott, K.W., and Rulten, S.L. (2014). One ring to bring them all-The role of Ku in mammalian non-homologous end joining. DNA Repair (Amst) 17. 10.1016/j.dnarep.2014.02.019.

9. Anderson, C.W., and Lees-Miller, S.P. (1992). The nuclear serine/threonine protein kinase DNA-PK. Crit Rev Eukaryot Gene Expr 2.

10. Grundy, G.J., Rulten, S.L., Arribas-Bosacoma, R., Davidson, K., Kozik, Z., Oliver, A.W., Pearl, L.H., and Caldecott, K.W. (2016). The Ku-binding motif is a conserved module for recruitment and stimulation of non-homologous end-joining proteins. Nat Commun 7. 10.1038/ncomms11242.

11. 11. Seif-El-Dahan, M., Kefala-Stavridi, A., Frit, P., Hardwick, S.W., Chirgadze, D.Y., Maia De Oliviera, T., Britton, S., Barboule, N., Bossaert, M., Pandurangan, A.P., et al. (2023). PAXX binding to the NHEJ machinery explains functional redundancy with XLF. Sci Adv 9. 10.1126/sciadv.adg2834.

12. Chen, S., Vogt, A., Lee, L., Naila, T., McKeown, R., Tomkinson, A.E., Lees-Miller, S.P., and He, Y. (2023). Cryo-EM visualization of DNA-PKcs structural intermediates in NHEJ. Sci Adv 9, eadg2838. 10.1126/sciadv.adg2838.

13. Chen, S., Lee, L., Naila, T., Fishbain, S., Wang, A., Tomkinson, A.E., Lees-Miller, S.P., and He, Y. (2021). Structural basis of long-range to short-range synaptic transition in NHEJ. Nature 593. 10.1038/s41586-021-03458-7.

14. 14. Chaplin, A.K., Hardwick, S.W., Liang, S., Kefala Stavridi, A., Hnizda, A., Cooper, L.R., De Oliveira, T.M., Chirgadze, D.Y., and Blundell, T.L. (2021). Dimers of DNA-PK create a stage for DNA double-strand break repair. Nat Struct Mol Biol 28. 10.1038/s41594-020-00517-x.

15. 15. Chaplin, A.K., Hardwick, S.W., Stavridi, A.K., Buehl, C.J., Goff, N.J., Ropars, V., Liang, S., De Oliveira, T.M., Chirgadze, D.Y., Meek, K., et al. (2021). Cryo-EM of NHEJ supercomplexes provides insights into DNA repair. Mol Cell 81. 10.1016/j.molcel.2021.07.005.

16. Stinson, B.M., Moreno, A.T., Walter, J.C., and Loparo, J.J. (2020). A Mechanism to Minimize Errors during Non-homologous End Joining. Mol Cell 77. 10.1016/j.molcel.2019.11.018.

17. Graham, T.G.W., Walter, J.C., and Loparo, J.J. (2016). Two-Stage Synapsis of DNA Ends during Non-homologous End Joining. Mol Cell 61. 10.1016/j.molcel.2016.02.010.

18. Amin, H., Zahid, S., Hall, C., & Chaplin, A. K. (2024). Cold snapshots of DNA repair: Cryo-EM structures of DNA-PKcs and NHEJ machinery. In Progress in Biophysics and Molecular Biology (Vol. 186). 10.1016/j.pbiomolbio.2023.11.007

19. Buehl, C.J., Goff, N.J., Hardwick, S.W., Gellert, M., Blundell, T.L., Yang, W., Chaplin, A.K., and Meek, K. (2023). Two distinct long-range synaptic complexes promote different aspects of end processing prior to repair of DNA breaks by non-homologous end joining. Mol Cell 83. 10.1016/j.molcel.2023.01.012.

20. Heyza, J. R., Mikhova, M., & Schmidt, J. C. (2023). Live cell single-molecule imaging to study DNA repair in human cells. In DNA Repair (Vol. 129). 10.1016/j.dnarep.2023.103540

21. Kim, J. S., Krasieva, T. B., Kurumizaka, H., Chen, D. J., Taylor, A. M. R., & Yokomori, K. (2005). Independent and sequential recruitment of NHEJ and HR factors to DNA damage sites in mammalian cells. Journal of Cell Biology, 170(3). 10.1083/jcb.200411083

22. Martinez-Pastor, B., Silveira, G. G., Clarke, T. L., Chung, D., Gu, Y., Cosentino, C., Davidow, L. S., Mata, G., Hassanieh, S., Salsman, J., Ciccia, A., Bae, N., Bedford, M. T., Megias, D., Rubin, L. L., Efeyan, A., Dellaire, G., & Mostoslavsky, R. (2021). Assessing kinetics and recruitment of DNA repair factors using high content screens. Cell Reports, 37(13). 10.1016/j.celrep.2021.110176

23. Kong, X., Mohanty, S. K., Stephens, J., Heale, J. T., Gomez-Godinez, V., Shi, L. Z., Kim, J. S., Yokomori, K., & Berns, M. W. (2009). Comparative analysis of different laser systems to study cellular responses to DNA damage in mammalian cells. Nucleic Acids Research, 37(9). 10.1093/nar/gkp221

24. Mavragani, I. V., Nikitaki, Z., Kalospyros, S. A., & Georgakilas, A. G. (2019). Ionizing radiation and complex DNA damage: From prediction to detection challenges and biological significance. In Cancers (Vol. 11, Issue 11). 10.3390/cancers11111789

25. Aleksandrov, R., Dotchev, A., Poser, I., Krastev, D., Georgiev, G., Panova, G., Babukov, Y., Danovski, G., Dyankova, T., Hubatsch, L., Ivanova, A., Atemin, A., Nedelcheva-Veleva, M. N., Hasse, S., Sarov, M., Buchholz, F., Hyman, A. A., Grill, S. W., & Stoynov, S. S. (2018). Protein Dynamics in Complex DNA Lesions. Molecular Cell, 69(6). 10.1016/j.molcel.2018.02.016

26. Heyza, J.R., Mikhova, M., Bahl, A., Broadbent, D., and Schmidt, J.C. (2023). Systematic analysis of the molecular and biophysical properties of key DNA damage response factors. Elife 12. 10.7554/eLife.87086.

27. Liu, P., Gan, W., Guo, C., Xie, A., Gao, D., Guo, J., Zhang, J., Willis, N., Su, A., Asara, J. M., Scully, R., & Wei, W. (2015). Akt-mediated phosphorylation of XLF impairs non-homologous end-joining DNA repair. Molecular Cell, 57(4). 10.1016/j.molcel.2015.01.005

28. 28. Schellenbauer, A., Guilly, M.N., Grall, R., Le Bars, R., Paget, V., Kortulewski, T., Sutcu, H., Mathé, C., Hullo, M., Biard, D., et al. (2021). Phospho-Ku70 induced by DNA damage interacts with RNA Pol II and promotes the formation of phospho-53BP1 foci to ensure optimal cNHEJ. Nucleic Acids Res 49. 10.1093/nar/gkab980.

29. Bekker-Jensen, S., Lukas, C., Kitagawa, R., Melander, F., Kastan, M.B., Bartek, J., and Lukas, J. (2006). Spatial organization of the mammalian genome surveillance machinery in response to DNA strand breaks. Journal of Cell Biology 173. 10.1083/jcb.200510130.

30. Mimori, T., Hardin, J.A., and Steitz, J.A. (1986). Characterization of the DNA-binding protein antigen Ku recognized by autoantibodies from patients with rheumatic disorders. Journal of Biological Chemistry 261, 2274–2278. 10.1016/S0021-9258(17)35929-X.

31. Yano, K., Morotomi-Yano, K., Wang, S., Uematsu, N., Lee, K., Asaithamby, A., Weterings, E., and Chen, D.J. (2008). Ku recruits XLF to DNA double-strand breaks. EMBO Rep 9, 91–96. 10.1038/sj.embor.7401137.

32. Hansen, A.S., Woringer, M., Grimm, J.B., Lavis, L.D., Tjian, R., and Darzacq, X. (2018). Robust model-based analysis of single-particle tracking experiments with spot-on. Elife 7. 10.7554/eLife.33125.

33. Tokunaga, M., Imamoto, N., and Sakata-Sogawa, K. (2008). Highly inclined thin illumination enables clear single-molecule imaging in cells. Nat Methods 5, 159–161. 10.1038/nmeth1171.

34. Koch, B., Sanchez, S., Schmidt, C. K., Swiersy, A., Jackson, S. P., & Schmidt, O. G. (2014). Confinement and Deformation of Single Cells and Their Nuclei Inside Size-Adapted Microtubes. Advanced Healthcare Materials, 3(11). 10.1002/adhm.201300678

35. Dedon, P.C., and Goldberg, I.H. (1992). Free-radical mechanisms involved in the formation of sequence-dependent bistranded DNA lesions by the antitumor antibiotics bleomycin, neocarzinostatin, and calicheamicin. Chem Res Toxicol 5, 311–332. 10.1021/tx00027a001.

36. Menon, V., and Povirk, L.F. (2016). End-processing nucleases and phosphodiesterases: An elite supporting cast for the non-homologous end joining pathway of DNA double-strand break repair. DNA Repair (Amst) 43, 57–68. 10.1016/j.dnarep.2016.05.011.

37. Leahy, J.J.J., Golding, B.T., Griffin, R.J., Hardcastle, I.R., Richardson, C., Rigoreau, L., and Smith, G.C.M. (2004). Identification of a highly potent and selective DNA-dependent protein kinase (DNA-PK) inhibitor (NU7441) by screening of chromenone libraries. Bioorg Med Chem Lett 14, 6083–6087. 10.1016/j.bmcl.2004.09.060.

38. Cui, X., Yu, Y., Gupta, S., Cho, Y.-M., Lees-Miller, S.P., and Meek, K. (2005). Autophosphorylation of DNA-dependent protein kinase regulates DNA end processing and may also alter double-strand break repair pathway choice. Mol Cell Biol 25, 10842–10852. 10.1128/MCB.25.24.10842-10852.2005.

39. Ding, Q., Reddy, Y.V.R., Wang, W., Woods, T., Douglas, P., Ramsden, D.A., Lees-Miller, S.P., and Meek, K. (2003). Autophosphorylation of the catalytic subunit of the DNA-dependent protein kinase is required for efficient end processing during DNA double-strand break repair. Mol Cell Biol 23, 5836–5848. 10.1128/MCB.23.16.5836-5848.2003.

40. Goodarzi, A.A., Yu, Y., Riballo, E., Douglas, P., Walker, S.A., Ye, R., Härer, C., Marchetti, C., Morrice, N., Jeggo, P.A., et al. (2006). DNA-PK autophosphorylation facilitates Artemis endonuclease activity. EMBO J 25, 3880–3889. 10.1038/sj.emboj.7601255.

41. Chan, D.W., and Lees-Miller, S.P. (1996). The DNA-dependent protein kinase is inactivated by autophosphorylation of the catalytic subunit. J Biol Chem 271, 8936–8941. 10.1074/jbc.271.15.8936.

42. Reddy, Y.V.R., Ding, Q., Lees-Miller, S.P., Meek, K., and Ramsden, D.A. (2004). Non-homologous end joining requires that the DNA-PK complex undergo an autophosphorylation-dependent rearrangement at DNA ends. J Biol Chem 279, 39408–39413. 10.1074/jbc.M406432200.

43. Block, W.D., Yu, Y., Merkle, D., Gifford, J.L., Ding, Q., Meek, K., and Lees-Miller, S.P. (2004). Autophosphorylation-dependent remodeling of the DNA-dependent protein kinase catalytic subunit regulates ligation of DNA ends. Nucleic Acids Res 32, 4351–4357. 10.1093/nar/gkh761.

44. 44. Goff, N. J., Brenière, M., Buehl, C. J., de Melo, A. J., Huskova, H., Ochi, T., Blundell, T. L., Mao, W., Yu, K., Modesti, M., & Meek, K. (2022). Catalytically inactive DNA ligase IV promotes DNA repair in living cells. Nucleic Acids Research, 50(19). 10.1093/nar/gkac913

45. Jayaram, S., Ketner, G., Adachi, N., & Hanakahi, L. A. (2008). Loss of DNA ligase IV prevents recognition of DNA by double-strand break repair proteins XRCC4 and XLF. Nucleic Acids Research, 36(18). 10.1093/nar/gkn552

46. Francis, D. B., Kozlov, M., Chavez, J., Chu, J., Malu, S., Hanna, M., & Cortes, P. (2014). DNA Ligase IV regulates XRCC4 nuclear localization. DNA Repair, 21. 10.1016/j.dnarep.2014.05.010

47. Huang, R. X., & Zhou, P. K. (2020). DNA damage response signaling pathways and targets for radiotherapy sensitization in cancer. In Signal Transduction and Targeted Therapy (Vol. 5, Issue 1). 10.1038/s41392-020-0150-x

48. Groelly, F. J., Fawkes, M., Dagg, R. A., Blackford, A. N., & Tarsounas, M. (2023). Targeting DNA damage response pathways in cancer. In Nature Reviews Cancer (Vol. 23, Issue 2). 10.1038/s41568-022-00535-5

49. Cong, L., Ran, F.A., Cox, D., Lin, S., Barretto, R., Habib, N., Hsu, P.D., Wu, X., Jiang, W., Marraffini, L.A., et al. (2013). Multiplex Genome Engineering Using CRISPR/Cas Systems. Science (1979) 339, 819–823. 10.1126/science.1231143.

50. Xi, L., Schmidt, J.C., Zaug, A.J., Ascarrunz, D.R., and Cech, T.R. (2015). A novel two-step genome editing strategy with CRISPR-Cas9 provides new insights into telomerase action and TERT gene expression. Genome Biol 16, 231. 10.1186/s13059-015-0791-1.

51. Hoelzel, C. A., & Zhang, X. (2020). Visualizing and Manipulating Biological Processes by Using HaloTag and SNAP-Tag Technologies. In ChemBioChem (Vol. 21, Issue 14). 10.1002/cbic.202000037

52. Zhao, C., Teng, E.M., Summers, R.G., Ming, G., and Gage, F.H. (2006). Distinct Morphological Stages of Dentate Granule Neuron Maturation in the Adult Mouse Hippocampus. The Journal of Neuroscience 26, 3–11. 10.1523/JNEUROSCI.3648-05.2006.

53. Neal, J.A., Xu, Y., Abe, M., Hendrickson, E., and Meek, K. (2016). Restoration of ATM Expression in DNA-PKcs-Deficient Cells Inhibits Signal End Joining. J Immunol 196, 3032–3042. 10.4049/jimmunol.1501654.

54. Broadbent, D.G., Barnaba, C., Perez, G.I., and Schmidt, J.C. (2023). Quantitative analysis of autophagy reveals the role of ATG9 and ATG2 in autophagosome formation. Journal of Cell Biology 222. 10.1083/jcb.202210078.

55. Laine, R.F., Tosheva, K.L., Gustafsson, N., Gray, R.D.M., Almada, P., Albrecht, D., Risa, G.T., Hurtig, F., Lindås, A.C., Baum, B., et al. (2019). NanoJ: A high-performance open-source super-resolution microscopy toolbox. J Phys D Appl Phys 52. 10.1088/1361-6463/ab0261.

56. Grimm, J.B., English, B.P., Chen, J., Slaughter, J.P., Zhang, Z., Revyakin, A., Patel, R., Macklin, J.J., Normanno, D., Singer, R.H., et al. (2015). A general method to improve fluorophores for live-cell and single-molecule microscopy. Nat Methods 12. 10.1038/nmeth.3256.

57. Grimm, J.B., Xie, L., Casler, J.C., Patel, R., Tkachuk, A.N., Falco, N., Choi, H., Lippincott-Schwartz, J., Brown, T.A., Glick, B.S., et al. (2021). A General Method to Improve Fluorophores Using Deuterated Auxochromes. JACS Au 1. 10.1021/jacsau.1c00006.

58. Klump, B.M., Perez, G.I., Patrick, E.M., Adams-Boone, K., Cohen, S.B., Han, L., Yu, K., and Schmidt, J.C. (2023). TCAB1 prevents nucleolar accumulation of the telomerase RNA to facilitate telomerase assembly. Cell Rep 42, 112577. 10.1016/j.celrep.2023.112577.

